# Enhancer-dependence of gene expression increases with developmental age

**DOI:** 10.1101/678334

**Authors:** Wenqing Cai, Jialiang Huang, Qian Zhu, Bin E. Li, Davide Seruggia, Pingzhu Zhou, Minh Nguyen, Yuko Fujiwara, Huafeng Xie, Zhenggang Yang, Danni Hong, Pengfei Ren, Jian Xu, William T. Pu, Guo-Cheng Yuan, Stuart H. Orkin

## Abstract

How overall principles of gene regulation (the “logic”) may change during ontogeny is largely unexplored. We compared transcriptomic, epigenomic and topological profiles in embryonic (EryP) and adult (EryD) erythroblasts. Despite reduced chromatin accessibility compared to EryP, distal chromatin of EryD is enriched in H3K27ac, Gata1 and Myb occupancy. In contrast to EryP-specific genes, which exhibit promoter-centric regulation through Gata1, EryD-specific genes employ distal enhancers for long-range regulation through enhancer-promoter looping, confirmed by Gata1 HiChIP. Genome editing demonstrated distal enhancers are required for gene expression in EryD but not in EryP. Applying a metric for enhancer-dependence of transcription, we observed a progressive reliance on enhancer control with increasing age of ontogeny among diverse primary cells and tissues of mouse and human origin. Our findings highlight fundamental and conserved differences in regulatory logic at distinct developmental stages, characterized by simpler promoter-centric regulation in embryonic cells and combinatorial enhancer-driven control in adult cells.

**Highlights:**

- Regulation of embryonic-specific erythroid genes is promoter-centric through Gata1
- Adult-specific control is combinatorial enhancer-driven and requires Myb
- Adult specific genes have increased enhancer-promoter chromatin interactions
- Enhancer-dependence increases progressively with increasing developmental age

## Introduction

Interactions between chromatin and nuclear regulatory factors establish gene expression programs during development (Long et al., 2016; Smale and Kadonaga, 2003). Whereas chromatin landscapes have been elucidated by genome-wide chromatin profiling methods in numerous adult cell types (Consortium, 2012; Kim et al., 2005), scant attention has been paid to embryonic cell types, other than embryonic stem (ES) cells. Whether the organization, or “logic”, of gene regulation differs between embryonic and adult type cells remains to be explored. We began by examining these issues in the context of functionally analogous blood cells of two different stages of ontogeny, and then extended our findings more broadly to other cell types.

Primitive (EryP; also referred to as “embryonic”) and definitive (EryD; also referred to as “adult”) erythroid cells constitute distinct, temporally overlapping lineages with similar *in vivo* function (i.e. oxygen transport and delivery in the circulation), and provide a unique opportunity to explore chromatin state at two different stages of ontogeny. Prior molecular studies have identified master erythroid-lineage transcription factors (TFs)--Gata1, Tal1 and Klf1-- and several adult-specific factors, namely Bcl11a, Sox6 and Myb (Cantor and Orkin, 2002; Mucenski et al., 1991; Palis, 2014; Sankaran et al., 2009; Xu et al., 2010). Whereas transcriptional mechanisms have been studied extensively in EryD cells, we are unaware of similar analyses in EryP cells, which arise in the yolk sac as a distinct lineage.

EryP cells emerge in blood islands, and mature as a semi-synchronous cohort in the circulation. EryD cells, which are adult-type, are generated first within the fetal liver and later in the postnatal bone marrow (Orkin and Zon, 2008). Gata1 plays a central role in the regulation of erythroid-specific genes in both EryP and EryD lineages and is required for their differentiation (Fujiwara et al., 1996; Pevny et al., 1995). Gata1 collaborates with other critical TFs, including Scl/Tal1, Ldb1, Fog1 and Lmo2 (Palis, 2014). Knockout of each of these genes leads to defects in EryP and EryD cells (Cantor et al., 2002; Cantor and Orkin, 2002; Mead et al., 2001; Palis, 2014). In contrast, the loss of Myb, which is expressed selectively in definitive type cells, impairs proliferation and differentiation of EryD, sparing EryP cells (Mucenski et al., 1991). Given the different reliance of EyP and EryD cells on TFs for their development, we have asked whether these related, but distinct cell lineages in ontogeny differ in their fundamental regulatory organization and logic.

To address this question, we isolated mouse EryP and EryD erythroblasts and characterized transcriptomes, chromatin accessibility, histone modifications, transcription factor (TF) occupancies, and 3D chromatin interactions. We observed that gene regulation in EryP is largely promoter-centric, whereas that in EryD was distal enhancer driven for activation. We hypothesized that these features reflect inherent differences between embryonic cells and more diverse, long-lived adult cell types. Analyses of available datasets of diverse mouse and human cells and tissues provided further support for the unexpected finding that regulatory logic changes with ontogeny.

## Results

### EryP and EryD transcription correlates differently with distal chromatin accessibility

CD71^+^/Ter119^+^ EryP and EryD cells were isolated by FACS from E10.5 embryonic peripheral blood and E13.5 fetal liver, respectively (Koulnis et al., 2011). We profiled transcriptomes, chromatin accessibility, histone modifications, TF occupancies and chromatin interactions (**Fig. 1A**). Identity of the respective cell populations was assessed by the expression of globin and TF genes characteristic of each lineage. RNA-seq and qRT-PCR confirmed expression of *Hbb-y* and *Hbb-bh2* in E10.5 CD71^+^/Ter119^+^ EryP cells (**Fig. S1A, S1B**), and *Hbb-b1* and *Hbb-b2* in E13.5 CD71^+^/Ter119^+^ EryD cells (**Fig. S1A, S1B**). Gata1 and Tal1 were highly expressed in EryP and EryD cells, whereas Bcl11a was expressed only in EryD cells (**Fig. S1A, S1C**). Comparative analysis of transcriptomes revealed 943 EryP-specific and 1,689 EryD-specific genes using DESeq2 (Love et al., 2014) (log_2_(fold-change)>1, *P*-value*<0.01*) (**Fig. 1B**). EryP-specific genes were modestly enriched with Gene Ontology (GO) terms associated with “metabolic process” (*P*=3.2E-5) and “pseudouridine synthesis” (*P*=4.0E-5), whereas EryD-specific expressed genes were significantly enriched in “cell cycle genes” (*P*=2.5E-12) and “erythrocyte development” (*P*=3.2E-8) (**Fig. 1C**). Consistent with the GO term “cell cycle genes” in EryD, human erythroid progenitors are produced at 2 million erythrocytes every second.

**Figure 1.**
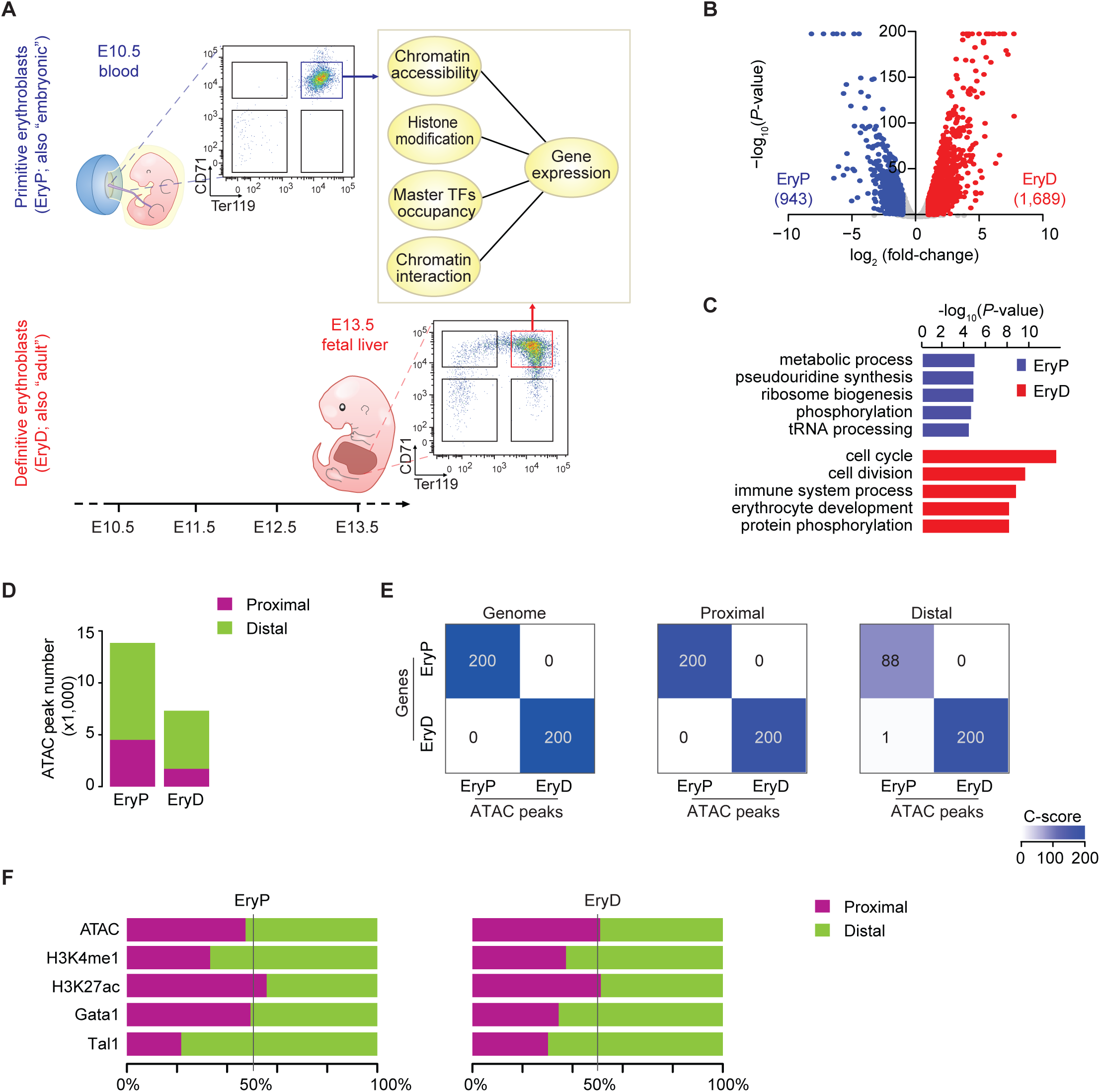
Chromatin accessibility differs in primitive and definitive erythroblasts. **(A)** Isolation of EryP and EryD and experiment outline. **(B)**Transcriptomic analysis of EryP and EryD. EryP-/EryD-specific expressed genes are highlighted in the volcano plot. **(C)** Gene Ontology (GO) terms enrichment in EryP-/EryD-specific expressed genes. **(D)** Comparisons of EryP-/EryD-specific ATAC peak numbers and their genomic distribution. **(E)** Association studies of EryP-/EryD-specific ATAC peaks and gene expression at genome-wide (left), proximal (middle) and distal (right) regions using C-score analysis. Rows are EryP-/EryD-specific genes. Columns are EryP-/EryD-specific ATAC peaks. C-score was defined as −log_10_ (*P*-value) using Fisher’s exact test (also see **Fig. S2A** and **Methods**), and C-scores ≥ 200 were presented as 200 to create appropriate color scale in heatmaps. **(F)** Genomic distribution of peaks of ATAC, histone marks and TFs. The graph shows the fraction of peaks at proximal and distal regions in EryP and EryD, respectively. See also Figure S1, S2 and Table S1, S2 and S6.

Nucleosome eviction due to regulatory factor binding is regarded as a primary step in transcriptional activation (Boyle et al., 2008; Neph et al., 2012b; Shlyueva et al., 2014; Thurman et al., 2012). To explore chromatin structure at the genome-wide level, we examined chromatin accessibility in EryP and EryD by Assay for Transposase-Accessible Chromatin using sequencing (ATAC-seq) (**Fig. 1D, 1F, Fig. S1E**). ATAC-seq retained high reproducibility in replicates (**Fig. S1D, S1G**). We assessed genome-wide differential accessibility between EryP and EryD using MAnorm (Huang et al., 2016; Shao et al., 2012; Xu et al., 2012) and observed 13,838 EryP-specific (4,506 at proximal and 9,332 at distal regions) and 7,315 EryD-specific (1,726 at proximal and 5,589 at distal regions) accessible regions (**Fig. 1D, Fig. S1F**), indicative of greater overall accessibility of EryP chromatin at both proximal and distal regions. A number of regions, such as the β-globin locus, displayed a consistent trend in terms of chromatin accessibility and gene expression in EryP and EryD (**Fig. S1G**).

To systematically assess the association between EryP- and EryD-specific accessible regions and differential gene expression, we introduced a computational metric, designated C-score (**Fig. S2A**), which we adopted from the Genomic Regions Enrichment of Annotations Tool (GREAT) analysis (McLean et al., 2010). GREAT predicts functions of cis-regulatory regions by analyzing the annotations of the nearby genes. To calculate a C-score, we thereby assigned EryP-/EryD-specific accessible regions (ATAC-seq peaks) to candidate genes using the ‘nearest neighbor gene’ approach (Huang et al., 2016; Xu et al., 2012). Then, the enrichment significance of a set of peak assigned EryP-/EryD-specific genes was assessed by using Fisher’s exact test, where −log_10_(*P*-value) was defined as the C-score (**Fig. S2A left**). Of note, C-scores increased as the association of accessible regions to specific expressed genes becomes greater. Within C-score analysis, EryP-/EryD-specific accessible regions, especially open proximal regions, were strongly correlated with EryP- and EryD-specific expressed genes (**Fig. 1E**), an observation consistent with the established association between chromatin accessibility and gene activation (Boyle et al., 2008; Neph et al., 2012b; Thurman et al., 2012). However, the C-score of distal accessible regions in EryD was greater than that in EryP (200 vs. 88) (**Fig. 1E right**), although we observed similar or even fewer distal differentially accessible regions in EryD as compared to EryP (5,589 vs. 9,332) (**Fig. 1D**). The divergence in the observed association likely reflects intrinsic differences in chromatin states within EryP- and EryD-specific distal accessible regions. Collectively, these data suggest that EryP- and EryD-specific distal accessible regions contribute differently to stage-specific gene expression.

### EryD-specific transcription is distal enhancer-driven

To evaluate whether the different association at distal regions in EryP and EryD relates to putative enhancer activity, we performed chromatin immunoprecipitation sequencing (ChIP-seq) for H3K27ac and H3K4me1. ChIP-seq of histone marks retained high reproducibility (**Fig. S1D, S1G**). As expected, the genome-wide distributions of the histone marks were consistent with published results (Wang et al., 2008). H3K4me1 was dominant at distal regions (**Fig. 1F, Fig.S1E**). H3K27ac, representing active chromatin, distributed across both proximal and distal regions (**Fig. 1F, Fig. S1E**). Next, we identified 5,137 EryP-specific H3K27ac peaks, 7,164 EryD-specific peaks and 11,234 shared peaks (**Fig. 2A, Fig. S1E, Fig. S3A**), and observed that the majority of cell-type-specific H3K27ac peaks was located at distal regions (**Fig. 2A**), which is consistent with the prevailing view that enhancers are more cell-type specific than promoters (Bulger and Groudine, 2010). Of note, while the numbers of cell-type specific H3K27ac peaks at proximal regions were comparable between EryD and EryP (1,652 vs. 1,841), we observed more cell-type specific distal H3K27ac peaks in EryD than in EryP (5,512 vs. 3,296) (**Fig. 2A**), suggesting that distal H3K27ac may be more correlated with expression in EryD. We further categorized the distal accessible regions based on H3K27ac peaks into two subgroups, active (with H3K27ac) or open only (without H3K27ac). Only a small fraction (22%) of EryP-specific distal accessible regions were active, which was similar to that in shared distal accessible regions (25%) (**Fig. 2B, Fig. S3B**). In contrast, the percentage was much greater (69%) in EryD-specific distal accessible regions (**Fig. 2B**). Besides, 81% of EryD-specific accessible regions in EryP were marked with H3K4me1 lacking H3K27ac (**Fig. S3C**). These findings suggest that EryD distal accessible regions are relatively enriched for more active chromatin, presumably reflecting distal active enhancers.

**Figure 2.**
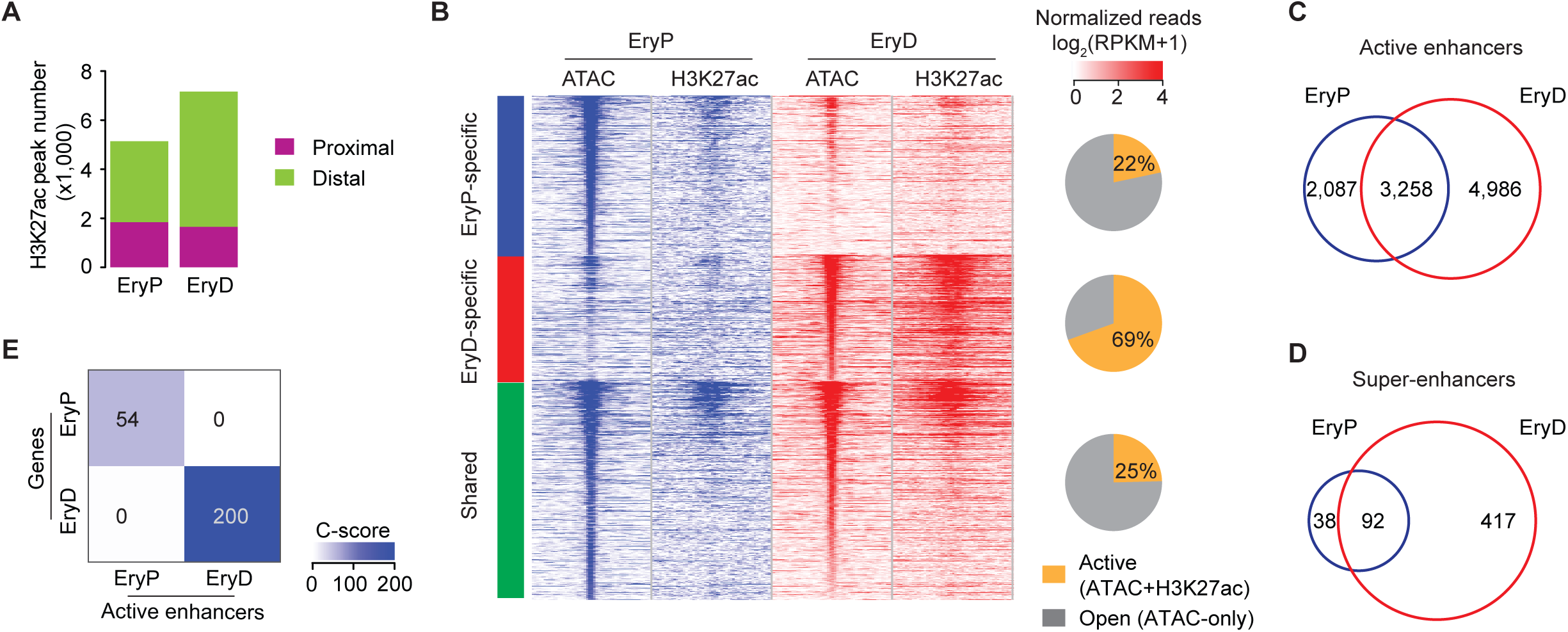
Chromatin state of distal accessible regions differs in primitive and definitive erythroblasts. **(A)** Comparisons of EryP-/EryD-specific H3K27ac ChIP-seq peak numbers and their genomic distribution. **(B)** Categorization of distal accessible regions with H3K27ac. Heatmaps of ATAC-seq and H3K27ac ChIP-seq signal, centered at the ATAC-seq peak summits. Chromatin distal accessible regions were grouped into three: EryP-/EryD-specific and shared peaks. Pie charts show the percentage of “active” (ATAC+H3K27ac) and “open” (ATAC-only) of each group, based on the presence of H3K27ac peaks. **(C-D)** Venn diagrams of EryP-/EryD-specific active enhancers (**C**) and super-enhancers (**D**). **(E)** Association study of EryP-/EryD-specific active enhancers and gene expression using C-score analysis. Rows are EryP-/EryD-specific genes. Columns are EryP-/EryD-specific enhancers. Also see **Fig. S2A** and **Methods**. See also Figure S1, S2, S3 and Table S1 and S2.

Distal enhancers are critical for tissue-specific gene expression patterns during lineage commitment. In total, we identified 2,087 EryP-specific, 4,986 EryD-specific and 3,258 shared active enhancers (**Fig. 2C**), which confirmed the greater overall number of H3K27ac peaks in EryD-specific distal regions. This relative difference became more extreme upon enumeration of “super-enhancers” (SEs), which were identified with the ROSE (Rank Ordering of Super-Enhancers) algorithm (Loven et al., 2013; Whyte et al., 2013). As shown in **Fig. 2D**, the number of EryD-specific SEs greatly exceeded that in EryP. GREAT analysis of EryP- and EryD-shared enhancers indeed revealed that most GO terms were associated with red blood cell functions, such as “porphyrin-containing compound metabolic process” (*P*=1.3E-14) and “erythrocyte homeostasis” (*P*=2.0E-13) (**Fig. S3D**), suggesting shared enhancers reflect red blood cell identity (Chung et al., 2012; Palis, 2014). Next, we performed motif analysis in three active enhancer categories (EryP-specific, EryD-specific and shared), and identified 17 significantly enriched motifs (**Fig. S3E**). The most enriched motifs corresponded to master TFs (TAL1, GATA1 and NFE2) in both EryP and EryD (**Fig. S3E**), a finding consistent with established roles of Tal1 and Gata1 at both stages of ontogeny (Fujiwara et al., 1996; Pevny et al., 1995; Porcher et al., 1996; Shivdasani et al., 1995) and the function of NFE2 in regulation of globin gene transcription (Shivdasani and Orkin, 1995).

We focused on EryP-/EryD-specific enhancers and performed the C-score analysis. We observed strong association between EryD-specific active enhancers and EryD-specific expressed genes, but weak association in EryP cells (**Fig. 2E**). To exclude false positives, we validated C-score analysis using different mapping approaches and alternative scoring metric (**Fig. S2A**, **Methods**). As shown in **Fig. S2A-S2D,** approaches of “multiple genes mapping” and “various mapping distance” achieved consistent C-score patterns as compared with the initial C-score analysis (**Fig. S2B-S2D, Fig. 2E, Methods**). Binomial test, which was used in GREAT analysis to assess functional significance of *cis-*regulatory elements across the entire genome (McLean et al., 2010), was adopted as an alternative scoring metric and −log_10_(Binomial *P*-value) was termed as GREAT score (G-score) (**Fig. S2A right**). In agreement with C-score analysis with EryP-/EryD-specific enhancers (**Fig. 2E**), we observed higher G-score in EryD-specific active enhancers and EryD-specific expressed genes than in EryP-specific enhancers and EryP-specific genes (**Fig. S2E**). These results indicate that conclusions derived from C-score analysis are reliable, and relatively insensitive to varying the above parameters. Taken together, these data suggest that transcription of EryD-specific genes is largely under the control of putative distal enhancers, whereas EryP-specific genes are more reliant on proximal elements.

### Gata1 controls EryP-specific gene transcription through proximal elements

To explore how master TFs and distinctive chromatin structures coordinately regulate transcription, we performed ChIP-seq analyses of Gata1 and Tal1 in both EryP and EryD (**Fig. 1F**). The total number of Tal1 lineage-specific peaks and their genomic distribution patterns were similar in EryP and EryD (**Fig. 3A, Fig. S3A**), and the majority of Tal1 peaks (>90%) resided at distal regions (**Fig. 3A**). EryD-specific Tal1 peaks were highly associated with EryD-specific expressed genes, whereas EryP-specific Tal1 peaks were weakly associated with EryP-specific expressed genes (**Fig. 3B, Fig. S4B**). This observation is consistent with higher enhancer-dependence of transcription in EryD than that in EryP (**Fig. 2E)**. In contrast to Tal1, Gata1 occupancy at distal regions was significantly greater in EryD than EryP (89% versus 57%), despite a comparable overall number of Gata1 peaks (**Fig. 3C, Fig. S3C**). In marked contrast, 43% Gata1 peaks in EryP were located in proximal regions (**Fig. 3C**), indicative of greater proximal Gata1 binding. We also observed strong association between EryP-specific Gata1 peaks and EryP-specific expressed genes (**Fig. 3D**), as contrasted with the pattern for Tal1 (**Fig. 3B)**. In particular, we observed that EryP-specific expressed genes were more enriched in EryP-specific Gata1 proximal peaks, whereas EryD-specific expressed genes were more enriched in EryD-specific Gata1 distal peaks (**Fig. 3E**). Consistent C-score patterns were also observed using different mapping approaches and alternative scoring metric in the C-score validation with Gata1 distal and proximal peaks (**Fig. S2F-S2I**). These data suggest that Gata1 regulates transcription in EryP predominantly through proximal regions. Upon comparison of Gata1 with H3K27ac profiles, we observed that Gata1 occupancy indeed co-localized with H3K27ac in both EryP and EryD (**Fig. 3G**). Taken together, we speculate that EryP-specific genes are regulated principally through Gata1 occupancy at proximal regions, whereas activation of EryD-specific genes is controlled largely by distal enhancers.

**Figure 3.**
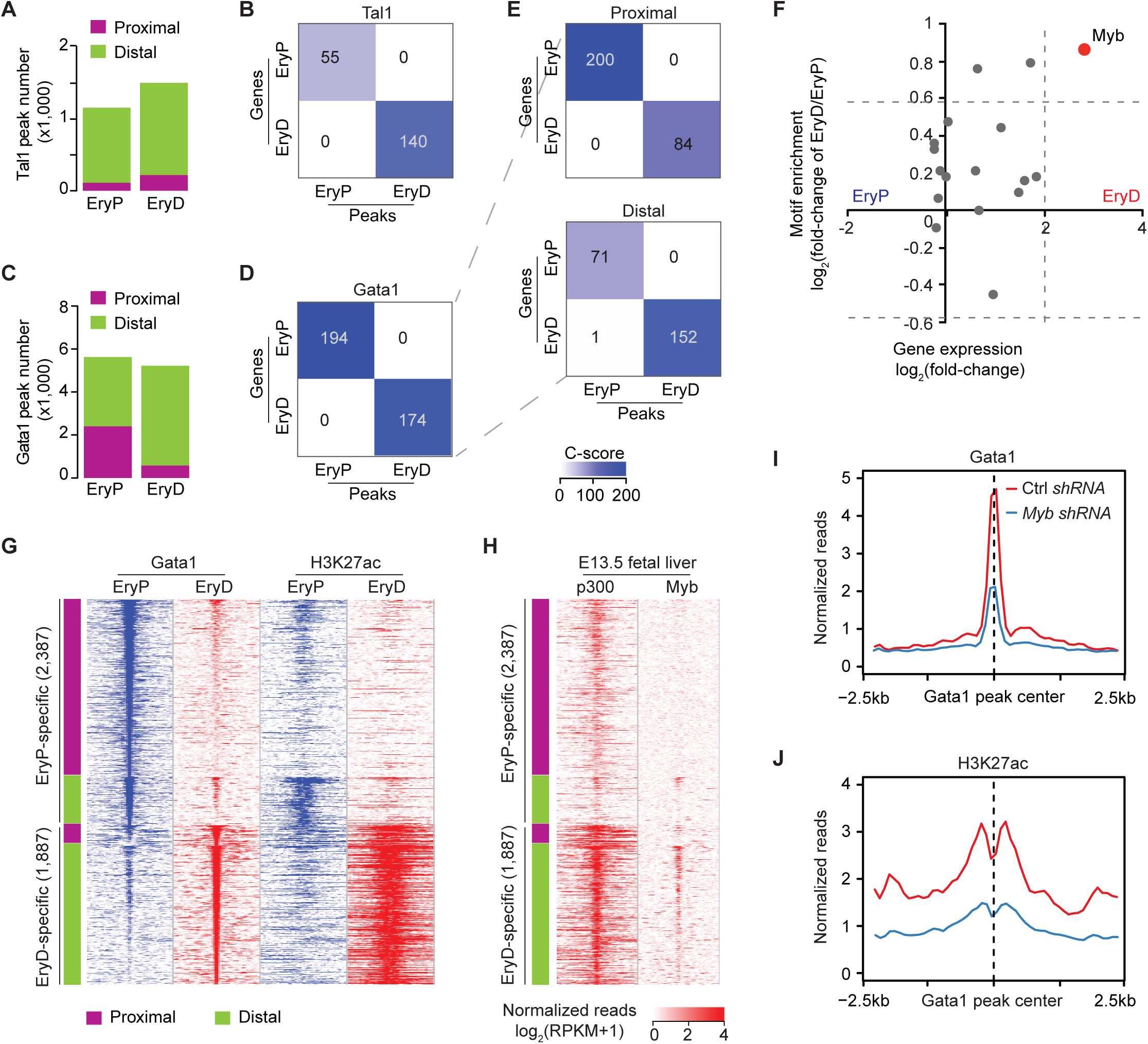
Myb is required for distal enhancer activation and Gata1 binding in definitive erythroblasts. **(A-B)** Comparison of EryP-/EryD-specific Tal1 peak numbers and their genomic distribution (**A**). Corresponding association study of EryP-/EryD-specific Tal1 peaks and gene expression is shown in (**B**). **(C-E)** Comparison of EryP-/EryD-specific Gata1 peaks and their genomic distribution (**C**). Corresponding association studies of EryP-/EryD-specific Gata1 peaks and gene expression are shown at genome-wide (**D**), proximal (**E, top**) and distal (**E, bottom**) regions. Also see **Fig. S2A** and **Fig.S2F-2I** for the validation of C-score analysis with Gata1 peaks. **(F)** Scatter plot of the difference of motif enrichment scores (*y*-axis) and gene expression (*x*-axis) reveals Myb (red spot) as an EryD-specific transcription factor. *y*-axis represents the log_2_ fold-change of the percentage of EryP-/EryD-specific enhancers with motifs, while *x*-axis represents the log_2_ fold-change of gene expression of the cognate TFs. 17 TF motifs were identified as significantly enriched in EryP- or EryD-specific active enhancers with *P*-value<1.0E-5 (**Fig. S3E**). **(G)** Heatmaps of the normalized reads of Gata1 ChIP-seq and H3K27ac ChIP-seq centered around the Gata1 peak summits in EryP and EryD. EryP-/EryD-specific Gata1 peaks were separated into proximal (purple) and distal (green) regions. **(H)** Heatmaps of the normalized reads of p300 and Myb ChIP-seq in E13.5 fetal liver, centered around Gata1 peak summits in EryP and EryD shown in Fig. 2G. **(I-J)** ChIP-seq signal density plots of Gata1 (**I**) and H3K27ac (**J**) in MEL cells treated with control *shRNA* (Ctrl *shRNA*) or *Myb shRNA*. Signals included in plots were restricted to EryD-specific distal regions as shown in Fig. 2G. *y*-axis is normalized ChIP-seq reads, log_2_(RPKM+1). MEL cells transduced with lentivirus carrying Dox-inducible *shRNA* directed to *Myb* or a control *shRNA*. Transduced cells were treated with 2 □g/ml Dox for 7 days, and harvested for H3K27ac and Gata1 ChIP-seq. See also Figure S2, S4 and Table S3.

### Myb mediates enhancer activation and Gata1 distal occupancy in EryD

We next sought to identify TFs that mediate EryD-specific enhancer activity and the distal distribution of Gata1 occupancy. Motif enrichment analysis in EryP-specific and EryD-specific enhancers suggested putative EryD-specific factors (**Fig. S3E**). We ranked TF motifs according to their relative enrichment in EryP-/EryD-specific enhancers, as well as expression of the cognate TFs in EryP and EryD (**Fig. 3F**). We found that Myb motif is more enriched in EryD-specific enhancers (**Fig. 3F**) and *Myb* is also specifically expressed in EryD (**Fig. 3F**). Previous work demonstrated that the loss of Myb impairs proliferation and differentiation of EryD (Mucenski et al., 1991), and the transcriptional coactivator CBP/p300, acetylating H3K27 (Tie et al., 2009), interacts with Myb through its KIX domain (Dai et al., 1996). In EryD cells, point mutation of the KIX domain phenocopies Myb loss (Kasper et al., 2002; Parker et al., 1999). Co-occupancy of Myb and CBP/p300 was also observed at super enhancers in T-cell acute lymphoblastic leukemia cells (Mansour et al., 2014). Moreover, expression of *p300* is higher in EryD than EryP (**Fig. S4D**). Therefore, we hypothesized that interaction of Myb and p300 may mediate activation of EryD-specific enhancers and promote Gata1 distal occupancy. To this end, we performed ChIP-seq of Myb and p300 in E13.5 fetal liver cells (**Fig. S4E**). Due to poor reproducibility with available p300 antibodies, we performed ChIP-seq with cells harvested from birA-expressing mice in which FLAG and *bio* epitopes were knocked into the C-terminus of the endogenous p300 allele (Zhou et al., 2017). Consistent with this hypothesis, Myb bound exclusively to distal regions and co-occupied with Gata1 in EryD, and the occupancy of p300 and Myb was highly co-localized with EryD-specific Gata1-bound enhancers (**Fig. 3G, 3H**) (*P*-value<2.2E-16). In addition, we examined effects of Myb loss of function. Since fetal erythropoiesis fails to occur in *Myb* null mice (Mucenski et al., 1991), we attenuated Myb expression in mouse erythroleukemia (MEL) cells with Doxycycline (Dox)-inducible *shRNA* directed to *Myb* or a control *shRNA*. Of note, MEL cells showed comparable chromatin landscape profiles as EryD (**Fig. 3G, S4F**). As shown in **Fig. 3I. 3J**, *shRNA* targeting of *Myb* decreased overall Gata1 binding and H3K27ac at EryD-specific Gata1 occupied distal regions (**Fig. 3I, 3J, Fig. S4G**). Taken together, these findings provide evidence that Myb is essential for EryD-specific enhancer activation and Gata1 occupancy at accessible distal regions.

### Gata1 HiChIP reveals increased enhancer-promoter interactions in EryD

ChIPseq analysis suggested striking differences in the chromatin landscapes of EryP and EryD, highlighted by elevated active enhancers at the EryD stage. We hypothesized that these findings reflect fundamental changes in the 3D chromatin organization. Thus, we examined enhancer-promoter (E-P) interactions as a functional readout of the distal enhancers and their association with gene expression in EryP and EryD cells by use of HiChIP (Mumbach et al., 2016; Mumbach et al., 2017). We performed Gata1 HiChIP to in EryP and EryD cells (**Fig. S5A**). Interaction matrices of EryP and EryD at progressively higher resolution revealed chromatin domains (**Fig. 4A**), as previously reported in high-resolution HiChIP analyses of other cell lines (Mumbach et al., 2017). Notably, Gata1 HiChIP in EryD exhibited more evident chromatin interactions than in EryP at 5-kb resolution (**Fig. 4A**), as well as a larger number of loops when zoomed in on a 400-kb loci (**Fig. 4B**). Overall, Gata1 HiChIP revealed 39,200 loops in EryP and 71,918 loops in EryD (**Fig. 4C**). We next characterized enhancer-promoter interactions of EryP-/EryD-specific genes in EryP and EryD cells by examining reads distribution within ±100 kb window from promoters. In contrast to sporadic enhancer-promoter interactions of EryP-specific genes in both EryP and EryD cells (**Fig. 4D left**), promoters of EryD-specific genes exhibited frequent interactions with surrounding enhancers in EryD cells, as compared with EryP cells (**Fig. 4D right**). In addition, we counted enhancer-promoter loops of EryP-/EryD-specific genes in EryP and EryD cells. Enhancer-promoter loops of EryP-specific genes did not reveal obvious differences between EryP and EryD (*P*=0.47), whereas enhancer-promoter loops of EryD-specific genes were significantly greater in number in EryD cells (*P*<0.01) (**Fig. 4E, Fig. S5C, S5D**). For example, in EryD-specific expressed loci (*Mgll* and *Abtb1*) (**Fig. S5B**), more enhancer-promoter loops were observed in EryD than in EryP (**Fig. 4B**). These observations provide additional evidence that EryP-specific gene activation employs promoter-centric regulatory logic, in which long-range enhance-promoter looping is not a prominent feature, whereas EryD-specific gene expression depends more heavily on enhancer-driven logic, in which frequent enhancer-promoter looping is observed. To test if enhancer contribution is greater overall in EryD cells, we counted enhancer-promoter loops in EryP- and EryD-common expressed genes (**Methods**), and observed that enhancer-promoter loops of common expressed genes in EryD cells were significantly greater in number in EryD cells than in EryP cells (*P*<0.01) (**Fig. 4E, Fig. S5C, S5D**). Taken together, these observations indicate that increased long-range interactions are associated with EryD-specific active enhancers

**Figure 4.**
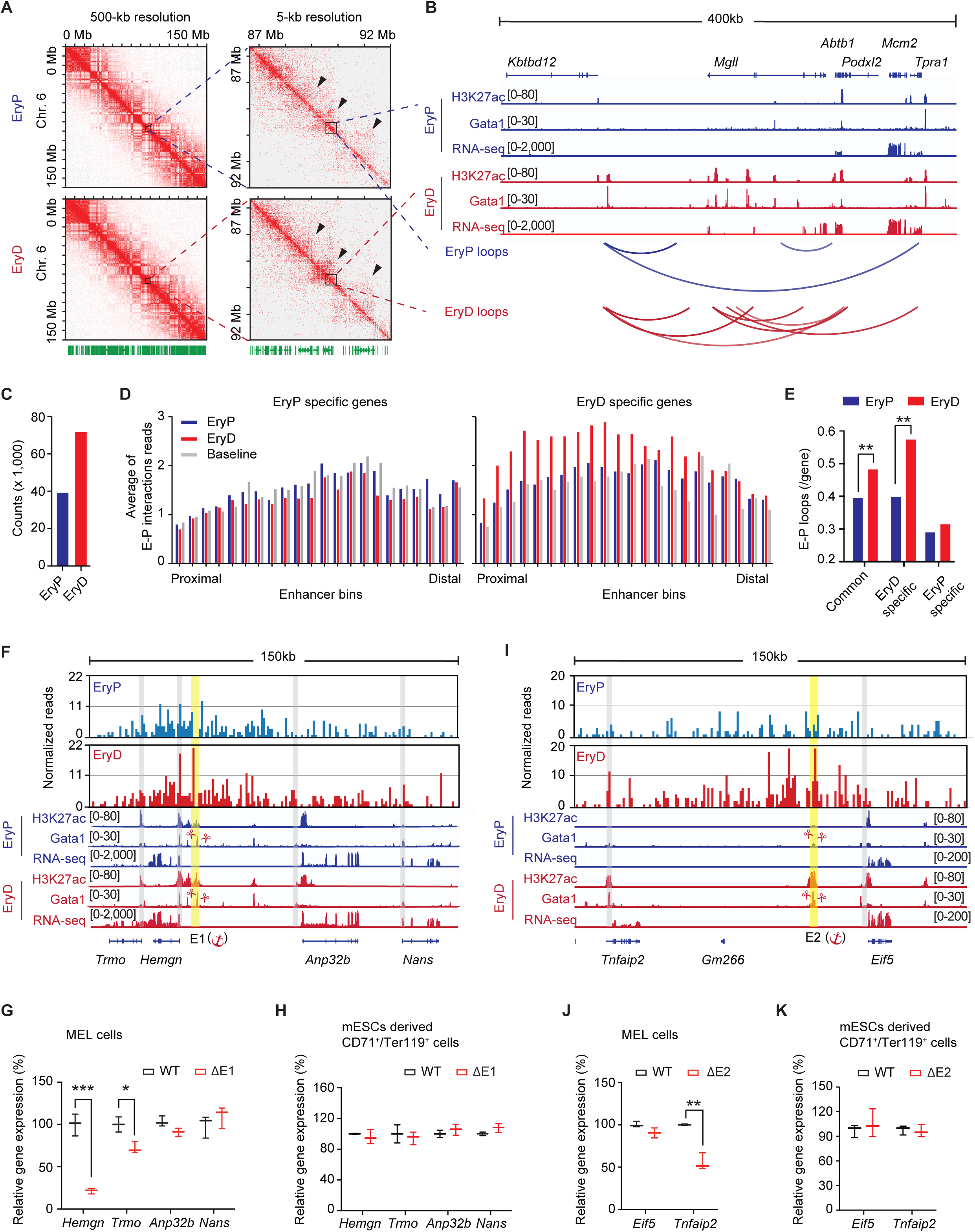
Distal regulatory elements play differential roles in primitive and definitive erythroblasts. **(A)** Interaction maps of Gata1 HiChIP in EryP and EryD at 500-kb and 5-kb resolution. Interaction maps were normalized by square root of vanilla coverage (VC SQRT). Zoom-in views of highlighted regions are in (**B**). **(B)** Gata1 HiChIP 3D signal enrichment at the *Mgll* and *Abtb1* loci in EryP and EryD cells. Representative plots of RNA-seq and ChIP-seq of H3K27ac and Gata1 from EryP and EryD cells were aligned to the genomic region. **(C)** Comparison of total loops in EryP and EryD. **(D)** Average profiles of enhancer-promoter (E-P) interactions of EryP-/EryD-specific genes. Promoters of EryP- or EryD-specific expressed genes were set as anchor points, and enhancers were defined by Gata1 peaks ±100 kb away from TSS. The average of reads within enhancer-promoter interactions of EryP- or EryD-specific expressed genes was plotted (see also **Methods**). *y*-axis is the average of normalized reads for E-P interactions per gene. Each enhancer bin in the *x*-axis indicates the rank position of the enhancer based on its distance to promoter. E-P interactions of EryP-/EryD-specific genes were comparaed with genomic baseline (grey bars), which are E-P interactions of a set of randomly selected genes of matched size in both EryP and EryD cells. **(E)** Quantification of enhancer-promoter (E-P) loops in EryP and EryD cells. Comparisons were at 25-kb resolution in three groups, EryP-specific genes, EryD-specific genes, and common expressed genes. *P*-value represents permutation test in 1,000 random genes selection of matched size. See also **Fig. S5C and S5D**. **(F)** Virtual 4C interaction profiles at the Enhancer 1 (E1) in the locus of *Trom, Hemgn, Anp32b and Nans* in EryP and EryD cells. Virtual 4C interaction profiles were generated at 1-kb resolution, and *y*-axis is normalized HiChIP reads. E1 is highlighted in yellow, and promoters of *Trom, Hemgn, Anp32b and Nans* are highlighted in grey. RNA-seq and ChIP-seq of H3K27ac and Gata1 from EryP and EryD cells were aligned to the genomic region. **(G-H)** Validation of E1 deletion in MEL cells (**G**) and mESCs-derived CD71^+^/Ter119^+^ cells (**H**). Quantitative transcript analysis of *Trom, Hemgn, Anp32b and Nans* upon E1 deletion in MEL cells (**G**) and mESCs-derived CD71^+^/Ter119^+^ cells (**H**). Single cell derived clones that had undergone biallelic excision of E1 were used in analyses (see also **Fig. S5J**). **(I)** Virtual 4C interaction profiles at the Enhancer 2 (E2) in the *Tnfaip2* and *Eif5* locus in EryP and EryD cells. E2 is highlighted in yellow, and promoters of *Tnfaip2* and *Eif5* are highlighted in grey. RNA-seq and ChIP-seq of H3K27ac and Gata1 from EryP and EryD cells were aligned to the genomic region. **(J-K)** Validation of E2 deletion in MEL cells (**J**) and mESCs-derived CD71^+^/Ter119^+^ cells (**K**). Quantitative transcript analysis of *Tnfaip2* and *Eif5* upon E2 deletion in MEL cells (**J**) and mESCs-derived CD71^+^/Ter119^+^ cells (**K**). Single cell derived clones that had undergone biallelic excision of E2 were used in analyses (see also **Fig. S5J**). ChIP-seq and RNA-seq track of EryP cells are labeled in blue and those of EryD cells are in red (**B**, **F** and **I**). Experiments were replicated at least twice in **G**, **H**, **J** and **K**. Error bars indicate the S.E.M.; n=3 (**G**, **H**, **J** and **K**). \**P*<0.05, \*\**P*<0.01, \*\*\**P*<0.001, unpaired one-tailed Student’s *t*-test (**G, H, J** and **K**) and permutation test (**E**). See also Figure S5 and Table S2, S4, S5 and S6.

### CRISPR/Cas9 genomic editing confirms putative enhancer-driven logic in EryD

We next disrupted specific enhancers to interrogate their requirement for gene expression in EryP and EryD cells. We focused on two EryD specific distal enhancers occupied by Gata1 (**Fig. 4F, 4I**): enhancer 1, or E1 (chr12: 111517811-111518523) in *Trmo*, *Hemgn*, *Anp32b* and *Nans* locus, and enhancer 2, or E2 (chr4: 46410632-46411247) in *Tnfaip2* and *Eif5* locus. *Trmo*, *Nans* and *Tnfaip2* are EryD-specific genes, whereas *Hemgn*, *Anp32b* and *Eif5* are EryP-/EryD common expressed genes (**Fig. S5E, S5F)**. We then applied Gata1 HiChIP to examine the 3D enhancer landscapes of the loci using virtual 4C (v4C) analysis (Mumbach et al., 2017). Here, we set the genomic position of a given enhancer as an anchor point (highlighted in yellow in **Fig. 4F** and **4I**) and visualized all interaction reads occurring with that anchor. As shown in the top two tracks (**Fig. 4F**), interaction reads between E1 and the *Hemgn* promoter (highlighted in grey) were greater in EryD cells than in EryP cells (27.2 in EryD *vs.* 19.7 in EryP at *Hemgn* promoter, and 22.2 in EryD *vs.* 8.7 in EryP at E1). Consistently, interaction reads between E2 and *Tnfaip2* promoter were greater in EryD than in EryP (15.5 in EryD *vs*. 1.8 in EryP at Tnfaip2 promoter, and 25.6 in EryD *vs*. 5.6 in EryP at E2) (**Fig. 4I**).

To evaluate enhancer contribution in EryD cells, we performed CRISPR/Cas9 in definitive stage (EryD) MEL cells (Ganguly and Skoultchi, 1985; Sheffery et al., 1984) using paired guide RNAs (sgRNAs) targeting 5’ and 3’ flanking regions of E1. The deletion size of the enhancer was refined by Gata1 ChIP-seq peaks (**Fig. 4F, 4I, Fig. S5J**). Transcripts of 4 genes in the E1 locus were examined in day 5 differentiated WT and ΔE1 MEL cells. *Hemgn* expression was profoundly reduced in the absence of the E1 enhancer (**Fig. 4G**). Additionally, *Trmo* transcripts were also significantly reduced in ΔE1 MEL cells (**Fig. 4G**). To evaluate enhancer contribution in EryP cells, we utilized mouse embryonic stem cells (mESCs) (Choi et al., 1998; Fraser et al., 2007; Zhang et al., 2003), as no primitive-stage erythroid cell lines are available. mESCs were differentiated into erythroblasts *in vitro* (**Fig. S5G, Methods**), and CD71^+^/Ter119^+^ cells were isolated by FACS for qRT-PCR analysis (Fraser et al., 2007)(**Fig. S5G**, **S5H**). Gene expression of *Hbb-b1*, *Hbb-bh1*, and *Hbb-by* in mESC-derived CD71^+^/Ter119^+^ cells indicated their primitive-stage origin (**Fig. S5I**) (Choi et al., 1998; Ganguly et al., 1985; Kingsley et al., 2013). In contrast to MEL cells, *Hemgn* and *Trmo* expression was maintained in enhancer deleted mESC-derived CD71^+^/Ter119^+^ cells (**Fig. 4H**). Of note, *Hemgn* is a EryP-/EryD-common expressed gene (**Fig. 4F, Fig. S5E**). Next, we used a similar strategy to validate E2 in MEL cells and mESC-derived CD71^+^/Ter119^+^ cells. Expression of *Tnfaip2* was markedly down-regulated in E2 deleted MEL cells (**Fig. 4J**), whereas *Tnfaip2* expression was unaffected in E2 deleted mESC-derived CD71^+^/Ter119^+^ cells (**Fig. 4K**). These findings provide clear evidence that distal enhancers are required for target gene expression at these loci in EryD, but not in EryP cells.

### Gene regulatory logic differ in EryP and EryD erythroblasts

The above analyses identified several features that distinguish EryP and EryD gene activation. We integrated data from ATAC-seq, ChIP-seq and RNA-seq and constructed gene regulatory association networks to present a coherent view of the differences between primitive and definitive red cell lineages (**Fig. 5A**). As depicted in **Fig. 5A**, the relative contribution of each feature (chromatin accessibility, TFs occupancy, and histone marks) to gene activation was assessed, and two gene activation modules (proximal and distal) have been identified for EryP and EryD, respectively. Proximal Gata1 peaks, co-localizing with ATAC-seq peaks, characterize EryP-specific gene expression (**Fig. 5A**). In contrast, a distal core-regulatory module in EryD, comprising multiple interactions of Gata1, H3K27ac and Myb, contributes to EryD-specific gene expression (**Fig. 5A**). Thus, EryP-specific gene expression is largely promoter-centric, whereas that in EryD cells reflects greater contribution of distal enhancers for gene activation (**Fig. 5A**).

**Figure 5.**
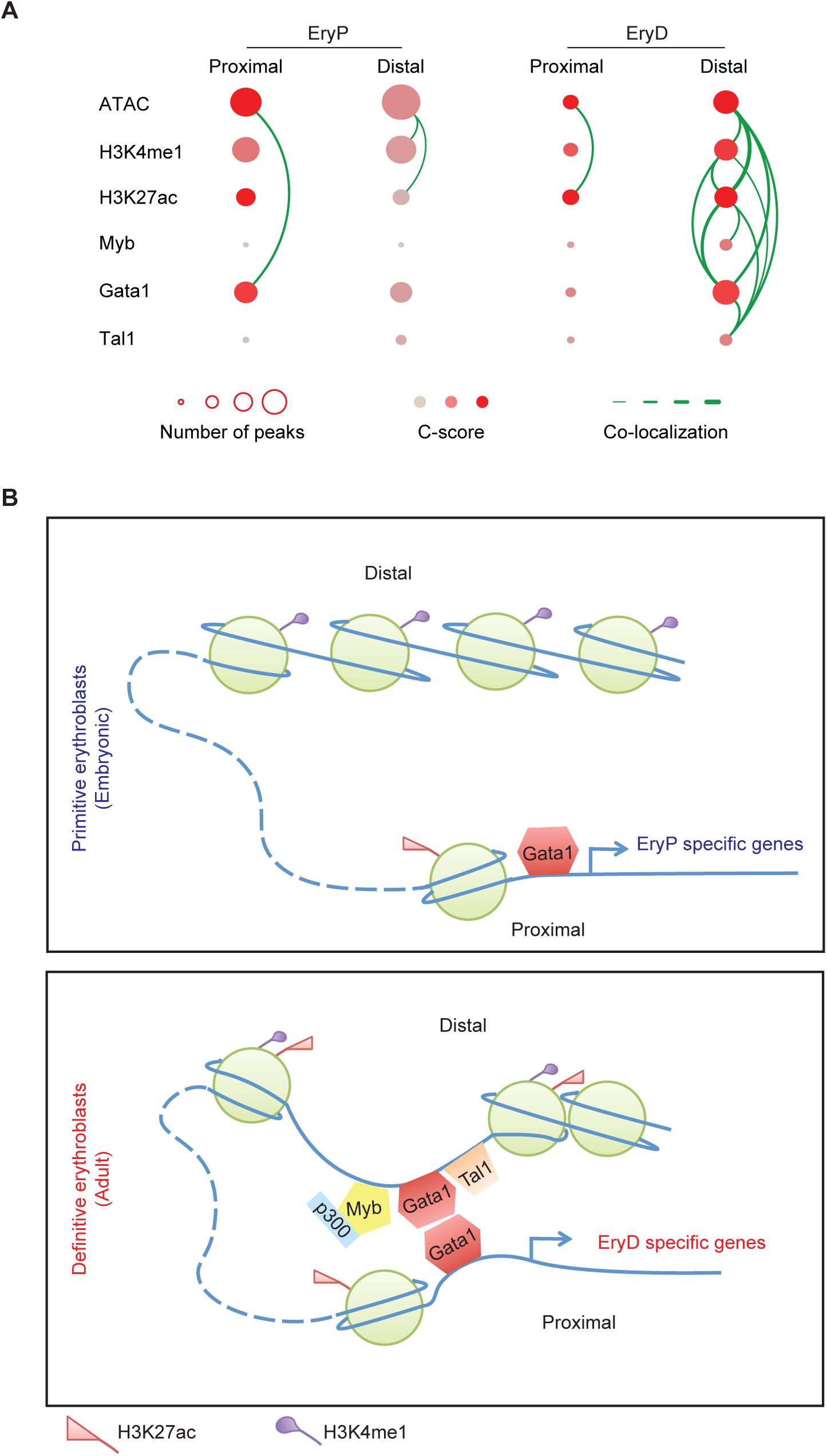
Distinct gene regulatory logic during the ontogeny of primitive and definitive erythroblasts. **(A)** The gene regulatory association networks in EryP and EryD cells. The association networks between cell-type-specific peaks and cell-type-specific expressed genes were constructed by integrating the information of ATAC-seq, ChIP-seq and RNA-seq data. Cell-type-specific peaks of ATAC-seq or ChIP-seq were divided into proximal and distal peaks based on their genomic distribution. The size of each node reflects the number of cell-type-specific peaks, while the node colors correspond to the C-score between cell-type-specific peaks and genes. Edges depict the −log_10_(*P*-value), where *P*-value represents the significance of peak co-localization, calculated using Fisher’s exact test. The threshold of 30 is used to determine whether an edge would be presented. (**B**) Summary of chromatin configuration during the ontogeny of erythropoiesis.

Taken together, genomic enhancer-promoter interaction studies and loss-of-function studies of selected enhancers further revealed a predominantly distal enhancer-driven regulatory logic in EryD, in contrast to promoter-centric regulatory logic in EryP (**Fig. 5B**). Promoter-centric regulatory logic in EryP is controlled principally through proximal Gata1, whereas distal enhancer-driven activation in EryD involves Myb and extensive enhancer-promoter interactions (**Fig. 5B**).

### Enhancer-driven regulatory logic correlates with development in mouse tissues

To explore whether our findings in primitive and definitive erythroid lineages are broadly relevant, we systematically evaluated the relative contribution of enhancer activities to gene regulation at different developmental stages by analyzing large-scale datasets generated by the mouse ENCODE consortium (Consortium, 2012) (**Fig. 6A, 6B, Methods**), using the C-score analysis described above. For these datasets, multiple embryonic tissues, including heart, limb, liver and neural systems, were collected anatomically, and the age was annotated as the number of days post coitum from E10.5 to E16.5 (**Fig. 6B**). In 42 qualified samples, we defined cell-type-specific genes (500 per cell type), in which the cell-type-specificity was calculated as the fold-enrichment of gene expression compared with the average of gene expression across all samples. The same strategy had been applied to define cell-type-specific enhancers (5000 per cell type), based on H3K27ac ChIP-seq signal. For each sample, the C-score was calculated by comparing the association between the sample-specific genes and sample-specific enhancers, as described above (**Fig. 6A, 6C**). As an additional control, we also applied the same method to evaluate the contribution of the enhancers specific to one sample to the expression pattern of a different sample. As expected, such cross-sample C-scores tend to have much lower values, especially if the samples are of different origins (**Fig. 6C, Fig. S6A**). The samples from same tissues or related tissues shown obvious modules in C-score matrix (**Fig. 6C, Fig. S6A**), indicating that C-scores reflect organ/tissue identity. Within each tissue, we observed a trend of increasing C-score within a given sample (diagonal values), correlating with progressively increased developmental age (**Fig. 6C**). To provide a statistical analysis of C-scores across developmental ages, we assigned samples of each tissue into three stages, “Early”, “Middle” and “Late”, based on their annotated information and dataset availability (**Fig. 6B, Methods**). In general, C-scores progressively increased with developmental age (**Fig. 6D**), and C-scores in the “Late” group were significantly higher than those in the “Early” group (**Fig. 6D**). Thus, progressive reliance of cell-specific gene expression on distal enhancers with developmental age apapears to be conserved during mouse development, in agreement with our findings in the erythroid lineages.

**Figure 6.**
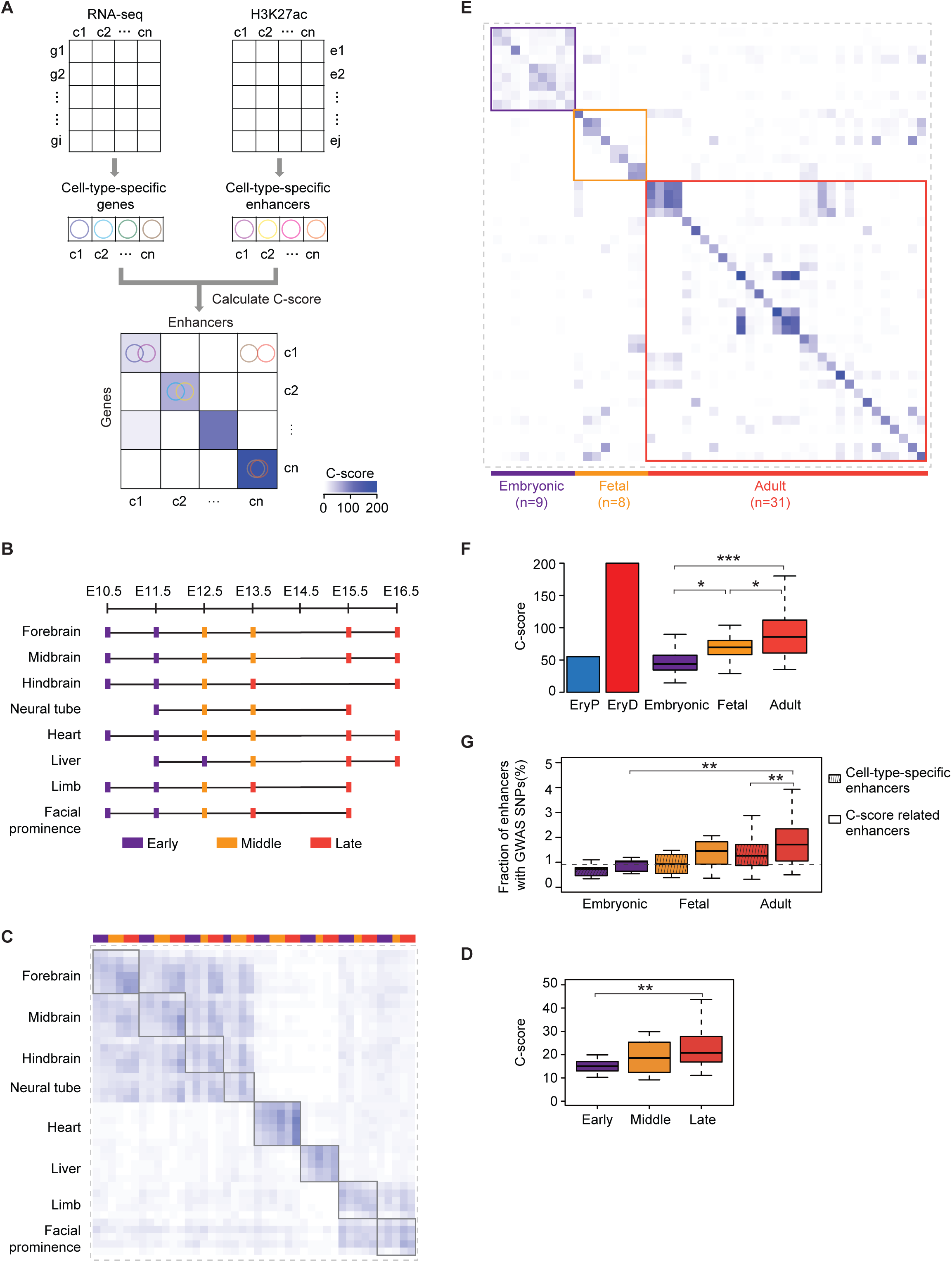
Enhancers contribute to cell-type-specific gene expression progressively with increasing age of ontogeny. **(A)**Schematic of unbiased evaluation of contribution of cell-type-specific enhancer to gene expression in ENCODE and Roadmap datasets. **(B)** The schematic diagram tissues and developmental ages in mouse ENCODE datasets. Samples from each tissue were classified into three stages as highlighted with different colors. **(C)** The C-score matrix from association studies of cell-type-specific enhancers and gene expression in mouse ENCODE datasets. Samples from the same tissue were aligned in a square box. Color bar next to *x*-axis indicates developmental stages in (**B**). Also see **Fig. S6A** for the C-score matrix with detailed annotation of tissues and developmental age. **(D)** Contribution of cell-type-specific enhancer to gene expression of ENCODE datasets, represented by the diagonal values in Fig. 6C, are summarized within different stages. **(E)** The C-score matrix from association studies of cell-type-specific enhancers and gene expression in Roadmap 48 human cell types, which were classified into three developmental stages, “Embryonic”, “Fetal” and “Adult”. See **Fig. S6B** for the C-score matrix with detailed annotation of cell types and C-scores. **(F)** Contribution of cell-type-specific enhancer to gene expression of Roadmap datasets, represented by the diagonal values in Fig. 6E, are summarized within different classes and compared with EryP and EryD. **(G)** Enrichment analysis of GWAS SNPs in cell-type-specific enhancers and C-score-related enhancers in Roadmap datasets. C-score-related enhancers are subset of cell-type-specific enhancers, in which nearby associated genes were also cell-type-specific expressed genes in C-score analyses. In box plot, the box represents the 25th and 75th percentiles, while the whiskers represent the 5th and 95th percentiles. \**P*<0.05, \*\**P*<0.01, \*\*\**P*<0.001, unpaired one-tailed Student’s *t*-test (**F, G,** and **H**). See also Figure S2 and S6.

### Enhancer-driven regulatory logic correlates with ontogeny in human cells

In order to provide an independent test of our hypothesis, we interrogated datasets of 48 diverse human cell types for which H3K27ac ChIP-seq and RNA-seq data are available from the NIH Roadmap project (Bernstein et al., 2010). The same analytical methods used to evaluate ENCODE datasets were applied to Roadmap datasets to define cell-type-specific genes and cell-type enhancers. Then, we performed C-score analysis and observed that the C-scores calculated within one cell type (diagonal values) were much higher than those across cell types (non-diagonal values) (**Fig. 6E, Fig. S6B**), consistent with the concept of tissue-specific enhancers (Andersson et al., 2014). Cell types were classified into three groups according to their developmental stage. An “Embryonic” group includes 9 ES cell lines (ESCs) or ES-derived cells. A “Fetal” group contains 8 primary cell types with annotation of fetal stage and up to 1 year old. An “Adult” group is comprised of 31 adult cell types isolated or derived from adult tissues (**Fig. S6B**). We observed that C-scores in the “Embryonic” group closely approximated that in EryP cells (**Fig. 6F**), whereas the overall correlation scores for the “Fetal” and “Adult” groups were significantly higher than those in the “Embryonic” group (**Fig. 6F**). Moreover, correlation scores increased progressively with developmental age (**Fig. 6F**). In addition, we performed the validation C-scores analysis for 48 human cell types (**Fig. S2A**), and observed that the pattern of C-score values among “Embryonic”, “Fetal” and “Adult” were consistent in all analyses (**Fig. S2K, S2L, S2M, S2N, Fig. 6F**). Recent studies suggest that disease-associated DNA sequence variation occurs largely in enhancers (Consortium, 2012; Hnisz et al., 2013). We next investigated the extent to which disease-associated SNPs occur in cell-type-specific enhancers in the three groups, and observed that “Adult” enhancers are highly enriched in GWAS SNPs (**Fig. 6G**). In addition, we focused on a subset of cell-type-specific enhancers, of which nearby associated genes were also cell-type-specific expressed in C-score analyses (**Fig. S2A**), termed as “C-score-related enhancers”. “C-score-related enhancers” were more strongly enriched with SNPs than cell-type-specific enhancers in the adult group (**Fig. 6G**), implying that variants might be associated with gene regulation and diseases. Among SNPs lying within C-score-related enhancers, rs755109 (chr9:100,696,202) (Lowe et al., 2009) is annotated to a cell-type-specific enhancer (chr9:100,695,253-100,696,742) and its nearby cell-type-specific expressed *HEMGN* in K562 cells and mobilized adult CD34^+^ primary cells. Using the lift-over tool of UCSC Genome Browser, we identified its conserved sequences, which coincidently overlapped the E1 enhancer (**Fig. 4F**). CRISPR/Cas9-mediated deletion of E1 led to decreased expression of *Hemgn* in mouse MEL cells but not in mESCs (**Fig. 4F, 4G** and **4I**). Taken together, these observations provide persuasive evidence that distinct contributions of promoter-centric and more combinatorial enhancer-driven regulation at embryonic and adult stages, respectively, represent a conserved theme through ontogeny.

## Discussion

Through comparative genome-wide analyses of embryonic and adult red cell lineages, we made the unanticipated observation that the dependence of cell-specific gene expression on distal enhancers increases with developmental stage. Embryonic erythroid gene expression is largely promoter-centric, whereas distal enhancers dominate regulatory control in adult-type cells. As these conclusions were largely inferred from the integration of cell-specific gene expression, chromatin accessibility, transcription factor binding, and active histone marks (e.g. H3K27ac), we employed HiChIP of the master erythroid transcription factor GATA1 and CRISPR/Cas9-mediated deletion of selected enhancers to test predictions from our initial conclusions. Indeed, we observed increased enhancer-promoter interactions in adult erythroid cells as compared with embryonic erythroid cells, and demonstrated a requirement for distal enhancers in adult, but not embryonic, cells. Taken together, our findings reveal that the extent to which gene expression relies on distal enhancers is not constant in development. As the vast majority of genome-wide analyses have focused on adult type cells, in which distal enhancers dominate transcriptional programs, our observations were unexpected.

Gene regulatory logic reflects the occupancy of *cis-*elements by transcription factors and the configuration of promoters and enhancers. Promoters, immediately upstream of transcriptional start sites, initiate transcription, whereas enhancers, located farther from genes, activate transcription through long-range looping, which has been revealed by ChIA-PET, HI-C and super-resolution microscopy (Krijger et al., 2016; Li et al., 2012; Schoenfelder and Fraser, 2019; Stadhouders et al., 2019). Tissue-specific enhancers, which ultimately determine tissue identity (Andersson et al., 2014), may reside at great distances (>1MB) from the gene body, and be brought in close proximity to promoters by looping (Schoenfelder and Fraser, 2019; Stadhouders et al., 2019). A distinguishing feature of our proposed regulatory networks of EryP and EryD is the relative contribution of proximal promoter-centric versus combinatorial enhancer control (**Fig. 5**). To ask whether our findings are relevant beyond the blood lineages we have studied, we performed computational analyses of available datasets of mouse and human origin encompassing tissues and cells of defined developmental age. In both species we observed progressive dependence of cell-specific gene expression on distal enhancers with developmental age (**Fig. 6D, 6F**). Thus, we have uncovered a conserved theme in development: regulatory logic is not invariant but instead changes in a progressive fashion to depend increasingly on distal enhancers.

Long-range communication between enhancers and promoters underlies complex gene regulatory networks (Bompadre and Andrey, 2019; Schoenfelder and Fraser, 2019; Stadhouders et al., 2019). Recent studies have shown that chromatin organization is reconfigured during stem cell differentiation and somatic cell reprogramming (de Laat and Duboule, 2013; Dixon et al., 2015; Krijger et al., 2016; Rao et al., 2014), suggesting stage specific chromating configuration instructs regulatory logic. Embryonic red cells represent a transient lineage, which is replaced by definitive red cells (adult lineage) that sustain the individual throughout life (Palis, 2014). Consistent with these notions, Gata1 HiChIP revealed a far greater number of enhancer-promoter loops of in EryD, as compared to embryonic EryP cells. Disruption, inversion or insertion of enhancers can perturb tissue-specific chromatin architecture and lead to inappropriate expression of target gene (Kragesteen et al., 2019; Kragesteen et al., 2018; Loven et al., 2013; Schuijers et al., 2018; Tickle and Towers, 2017; Williamson et al., 2016). Here, CRISPR/Cas9 mediated deletion of selected enhancers resulted in a striking decrease in target gene expression in adult stage MEL cells, but not in mESC-derived erythroblasts (**Fig. 4G, 4H, 4J, 4K**), suggesting the dependence of enhancers in mediating enhancer-promoter configuration is developmental age specific. The greater involvement of enhancers in long-range regulation in EryD requires activation of distal enhancers. We identified Myb as an EryD-specific regulator which exclusively bound to EryD-specific distal enhancers (**Fig. 3G-H**). Loss of *Myb* decreased overall H3K27ac deposition and Gata1 occupancy within these distal enhancer regions (**Fig. 3I-J**), suggesting that Myb is essential for EryD-specific distal enhancer activation. These observations imply that enhancer-promoter configuration is developmental age specific, involving specific transcription factors and enhancer activation.

The more dominant role of distal enhancers in adult differentiated cells may reflect their greater need to respond to complex and changing environmental cues, including cell-cell interactions, cytokines, soluble factors, and mechanic forces that trigger signal transduction to the nucleus (Schoenfelder and Fraser, 2019; Stadhouders et al., 2019). The shift of regulatory logic from promoter-centric in embryonic cells to enhancer-dependent in adult cells mirrors increased complexity of pathways and extracellular niches in the adult stage (Heinz et al., 2015). In contrast, the embryonic extracellular environment changes dynamically, for example at the onset of cardiac contractility at E9.0-E9.5 in the mouse embryo (Chen et al., 2014; Ji et al., 2003). Whatever the teleologic basis, promoter-centric regulation in embryonic cells and greater enhancer-dependent control in adult cells, respectively, constitute a previously unrecognized theme in network organization. The greater involvement of distal enhancers in adult lineages is relevant to human diseases, as the vast majority of disease-associated variations occur within non-coding genomic sequences (Consortium, 2012; Hnisz et al., 2013). Further in-depth genome-wide studies, including programmable 3D genome rewiring, may provide additional insights into the regulatory logic employed at different stages of ontogeny.

## Supporting information

All supplemental information

## Acknowledgments

We thank Dr. Jennifer Trowbridge at The Jackson Laboratory for assistance with ATAC-seq. We are grateful to Nicole Flanagan from the Center for Cancer Computational Biology sequencing facility, John Daley from the Flow Cytometry facility at DFCI, and Xiaoji Wu from HHMI sequencing facility, for technical help. We thank Drs. Partha Pratim Das, Sidinh Luc, Hye Ji Cha and Yan Kai for helpful discussions. Research was supported by funding from a NIH Cooperative Centers of Excellence in Hematology award (5U54DK11805) to S.H.O., NIH grants R01HL119099 and R01HG009663 to G.C.Y. and S.H.O., and the National Natural Science Foundation of China 31871317 to J.H. S.H.O. is an Investigator of the Howard Hughes Medical Institute.

## Author contributions

S.H.O. and G.C.Y. supervised this study. S.H.O., G.C.Y., W.C., and J.H. designed this study. W.C. and J.H. performed cell sorting, RNA-seq, ChIP-seq, and ATAC-seq. J.H., W.C., Z.Y., D.H. and P.R. performed computational analysis. W.C. and B.E.L. designed and performed Gata1 HiChIP. Q.Z. conducted HiChIP computational analysis. P.Z. designed and performed p300^fb^ bioChIP. W.C., M.N., and Y.F. performed experiments with embryos related work. W.C. and D.S. designed and conducted CRISPR/Cas9 genome editing in MEL cells and mESCs. J.X., H.X., and W.T.P. conceptually and technically provided feedbacks and helps. W.C., J.H., S.H.O. and G.C.Y. interpreted the results, and prepared manuscript with input from all authors.

## STAR Methods

### Key Resources Table

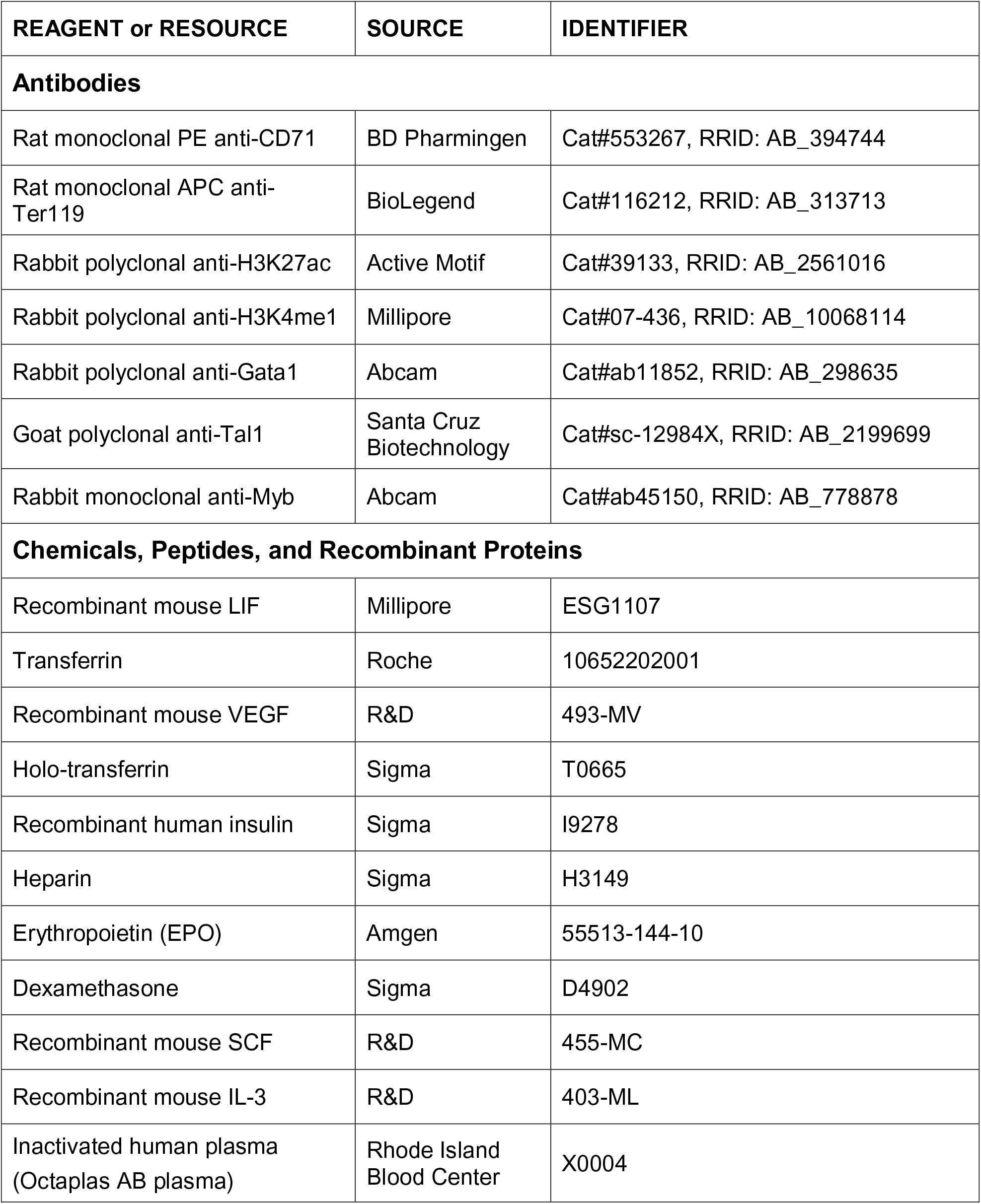
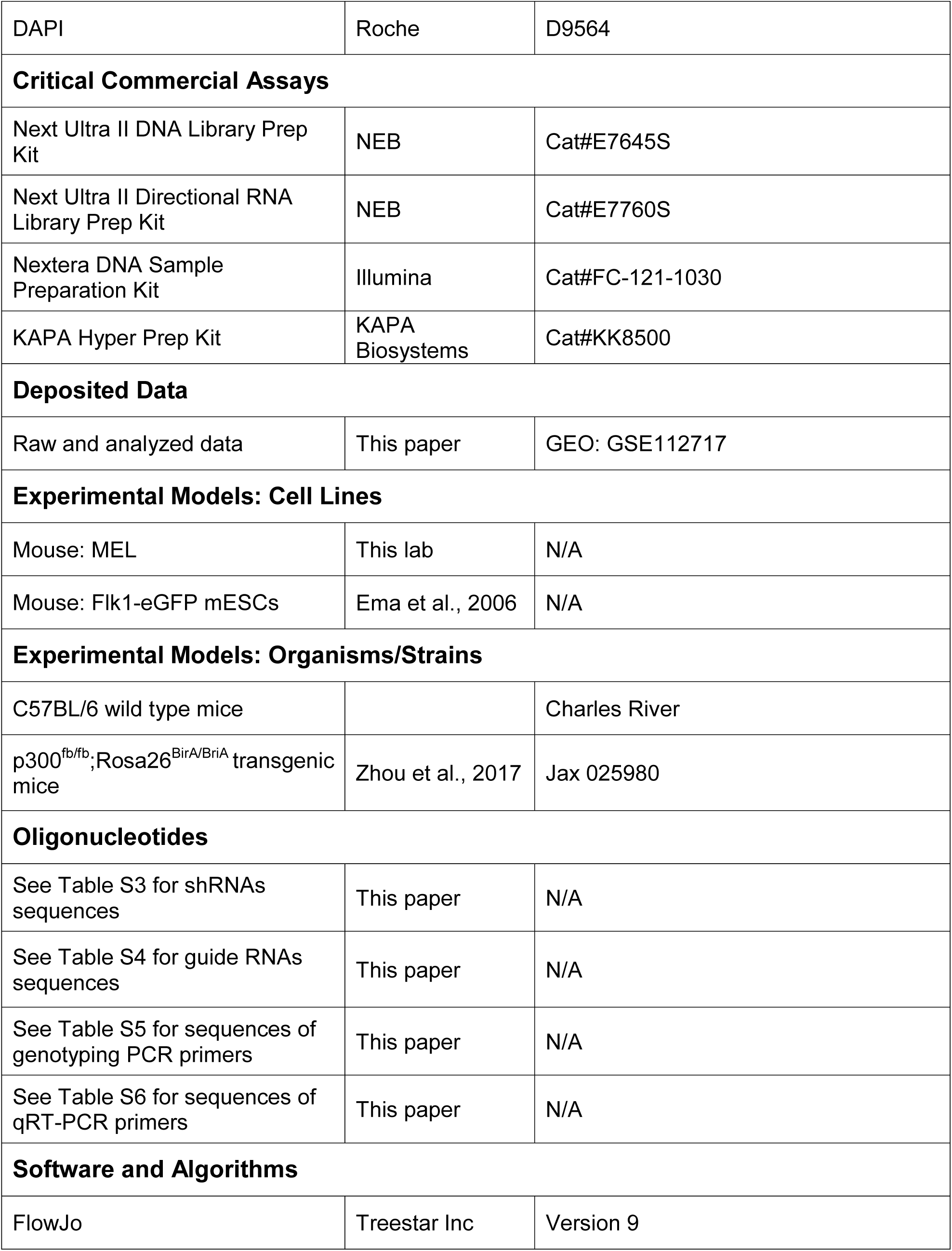
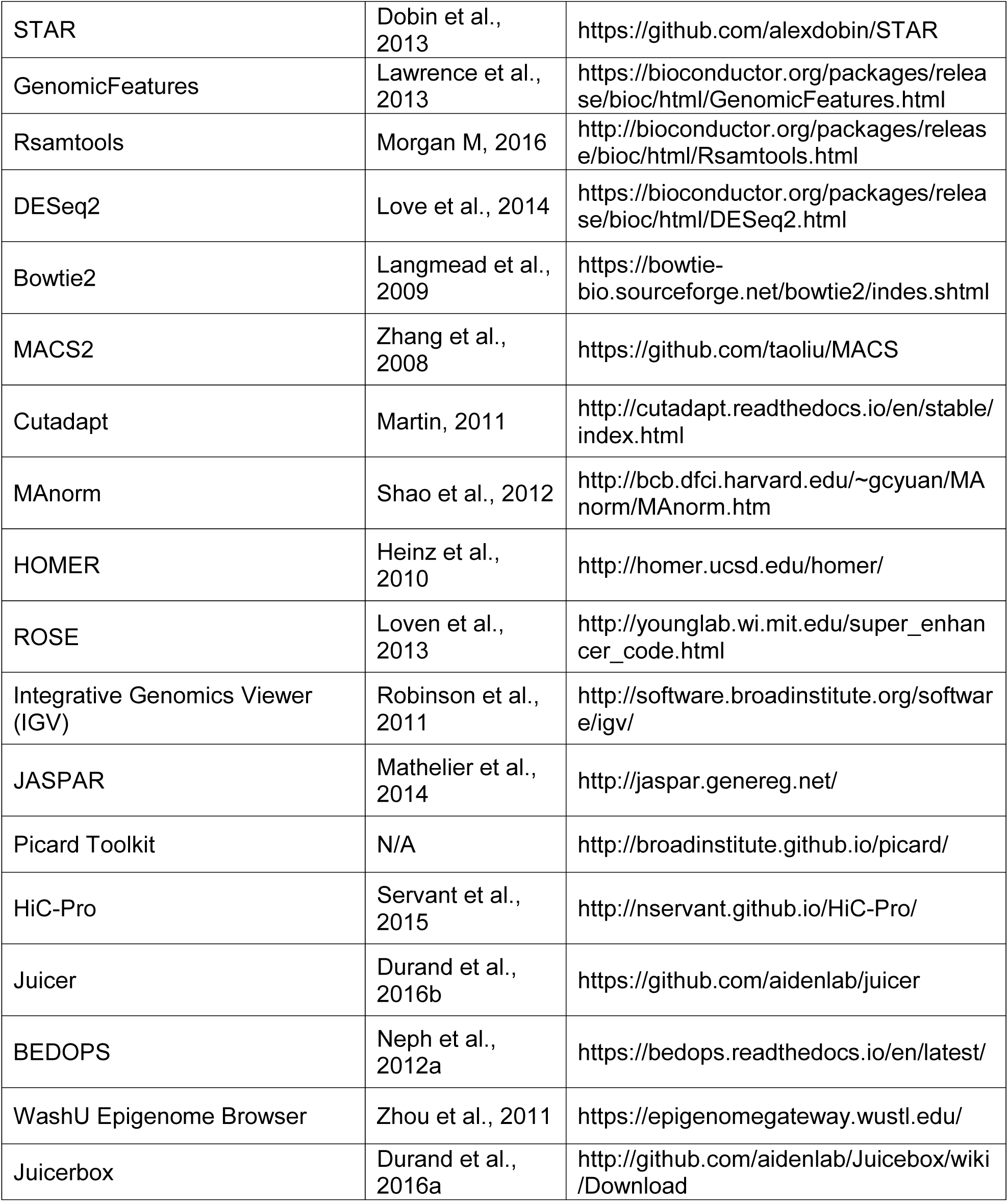

### Contact for Reagent and Resource Sharing

Further information and requests for reagents should be directed to, and will be fulfilled by the Lead Contact, Stuart H. Orkin.

### Experimental Model and Subject Details

#### Mice

E10.5 and E13.5 wild type mouse embryos were obtained from C57BL/6 females crossed to male mice. To perform p300^fb^ biochip, E13.5 fetal liver were obtained from Swiss Webster wild type females crossed to p300^fb/fb^;Rosa26^BirA/BriA^ males (Jax 025980).

#### Isolation of EryP and EryD erythroblasts by FACS

EryP were collected from peripheral blood of E10.5 mouse embryos in HBSS supplemented with 10% FBS and 0.25% EDTA. EryD were collected by mechanically dissociation of E13.5 whole fetal liver. Single cells were stained for CD71 (553267, BD Biosciences) and Ter119 (116212, BioLegend), and subsets were gated by the expression of CD71 and Ter119 as described (Koulnis et al., 2011). Live cells were sorted into CD71^+^/Ter119^+^ population by FACS. Dead cells were excluded using DAPI (Roche). Cell sorting was performed with J Ariall SORP UV (BD Biosciences) and analysis was performed on LSR II (BD Biosciences). FACS data were analyzed with FlowJo (Treestar Inc.).

#### MEL cell culture

MEL cells were cultured in DMEM supplemented with 10% fetal bovine serum, 2 mM *L*-glutamine and Penicillin and Streptomycin. 2% DMSO in culture medium was used to induce MEL cells differentiation. Medium was changed on differentiation day 3.

#### Mouse ESCs culture and differentiation

mESCs carrying Flk1-eGFP reporter (Ema et al., 2006) were cultured and differentiated as previously described (Cai et al., 2012). Briefly, mESCs were maintained with 1,000 unit/ml LIF (Millipore). In serum-induced differentiation, undifferentiated ESCs were dissociated into single cells by TrypLE (Thermo Fisher) and then differentiated as embryoid bodies (EBs) in Iscove’s Modified Dulbecco Media (IMDM) supplemented with 10% FBS, 2 mM *L*-Glutamine, 4.5×10^-4^ M monothioglycerol, 0.5 mM ascorbic acid, 200 μg/ml transferrin (Roche), 5% protein-free hybridoma media (PFHM-II; Life Technologies) and Penicillin and Streptomycin. Day 4 EBs were treated with 25 ng/ml recombinant VEGE (R&D Systems) for 3 days.

#### Primitive erythroid differentiation *in vitro*

To induce erythroid differentiation, day 7 EBs were dissociated into single cells by TrypLE (Thermo Fisher) and then cultured for 4 days in IMDM supplemented with 1% BSA, 200 μg/ml holo-transferrin (Sigma), 10 μg/ml recombinant human insulin (Sigma), 2 IU/ml Heparin (Sigma), 5% inactivated plasma (Rhode Island Blood Center), 2 mM *L*-Glutamine, 4.5×10^-4^ M β-mercaptoethanol, 3 U/ml Erythropoietin (EPO) (Amgen), 1 μM Dexamethasone (Sigma), 50 ng/ml recombinant mouse SCF (R&D), 10 ng/ml recombinant mouse IL-3 (R&D), and Penicillin and Streptomycin (Carotta et al., 2004; Zhang et al., 2003). Day 11 cultures were filter through 70 μM cell strainer to remove clumps, and then were cultured for another 4 days in above medium supplemented with 6 U/ml EPO (Amgen), 10 μM Dexamethasone (Sigma). Day 15 cultures were sorted into CD71^+^/Ter119^+^ population by FACS and followed by RNA extraction and qRT-PCR analysis. Dead cells were excluded using DAPI (Roche). Cell sorting was performed with J Ariall SORP UV (BD Biosciences).

### Method Detail

#### RNA-seq library preparation

Total RNA was extracted with TRIzol (Thermo Fisher) and cleaned with RNeasy Plus Mini Kit (QIAGEN) following manufacturer’s instructions. RNA-seq libraries were prepared using Next Ultra Directional RNA Library Prep Kit Illumina (New England Biolabs) according to the manufacturer’s instructions. Ilumina HiSeq 2500 was used for sequencing.

#### ATAC-seq library preparation

ATAC-seq was performed as described (Buenrostro et al., 2013) on FACS isolated EryP and EryD cells. 2.5 x 10^4^ cells were used per sample. 2.5 μl Tn5 Transposases (Illuminar) was used to make the transposition reaction. Libraries were generated using Ad1_noMX and Ad2.1-2.4 barcoded primers (Buenrostro et al., 2013) and were amplified for 6-9 total cycles. All libraries were sequenced on Illumina HiSeq2000 with 100bp paired-end reads.

#### Chromatin immunoprecipitation and ChIP-seq library preparation

ChIP was performed as previously described (Cai et al., 2013). FACS isolated cells, MEL cells or dissociated fetal liver cells were cross-linked with 1% v/v formaldehyde at room temperature for 10 minutes, and were quenched with 0.125 M Glycine for 5 minutes. Cells were washed twice in cold PBS, and cell pellets were stored at −80°C. Cell pellets were lysed in Nuclear Lysis Buffer (50 mM Tris-HCL pH 7.5, 5 mM EDTA, 0.1% SDS, 0.5% NP40, 150 mM NaCl, 0.25% Sarkosyl and 1 mM DTT) and followed by sonication to generate 150-300bp fragments using a Covaris E220 with following parameters: Duty Factor=5.0, Peak Power=140, Cycle/Burst=200, Time=900 seconds. Chromatin extracts were immunoprecipitated with antibodies overnight at 4°C, and then incubated with protein A Dynabeads (Invitrogen) for another 4 hours. 1.5 million cells were used for each Histone antibody, and 3-6 million cells were used for Tal1 and Gata1 antibody. 30 million unsorted fetal liver cells were used for Myb antibody. ChIP complexes were eluted and cross-linking reversed by heating at 65°C overnight. DNA was extracted with UltraPure Phenol:Chloroform:Isoamyl Alcohol (Thermo Fisher). The following antibodies were used: H3K27ac (39133, Active Motif), H3K4me1 (07-436, Millipore), Gata1 (ab11852, Abcam), Tal1 (sc-12984X, Santa Cruz), and Myb (ab45150, Abcam).

1 ng of ChIP DNA was used to prepare multiplexed sequencing libraries with Next Ultra II DNA Library Prep Kit for Illumina (New England BioLabs) according to the manufacturer’s instructions. Illumina HiSeq2500 or NextSeq 500 was used for sequencing.

#### p300^fb^ bioChip

E13.5 fetal livers were isolated from pregnant Swiss Webster wild type females crossed to p300^fb/fb^;Rosa26^BirA/BriA^ males (Jax 025980). Fetal liver cells were dissociated and then cross-linked in 1% v/v formaldehyde-containing at room temperature for 15 min. Glycine was added to final concentration of 0.125 M to quench formaldehyde. Cell pellets were lysed and followed by sonication to generate about 500bp fragments. For p300 bioChIP, the chromatin from 50 million cells was incubated with 100 µl Dynabeads M-280 Streptavidin (Life Technologies, 11206D) for 1 hour at 4°C. The streptavidin beads were washed and bound DNA eluted as previously described (Zhou et al., 2017). BioChIP DNA was purified with MinElute PCR Purification kit (Qiagen, 28006). ChIP-seq libraries were constructed using a ChIP-seq library preparation kit (KAPA Biosystems KK8500). 50 ng of sonicated chromatin without pull-down was used as input.

#### Gata1 HiChIP

HiChIP was performed as previously described (Mumbach et al., 2016) using the Gata1 antibody (ab11852, Abcam). In brief, 15 million of either peripheral blood cells of E10.5 mouse embryos or single cells of E14.5 fetal liver were cross-linked with 1% v/v formaldehyde at room temperature for 10 minutes, and were quenched with 0.125 M Glycine for 5 minutes. Crosslinked cells were lysed in cold Hi-C lysis buffer (10 mM Tris-HCl pH 7.5, 10 mM NaCl, 0.2% NP40, 1x Roche protease inhibitors) and followed by 1 U/μl MboI (NEB) digestion at 37°C for 6 hours, fill-in of overhangs with biotin-dATP (Thermo Fisher) at 37°C for 1.5 hours, and ligation with 4 U/μl T4 DNA Ligase (NEB) at room temperature for 6 hours. The nuclear pellet was sheared using a Covaris E220 with following parameters: Duty Factor=5.0, Peak Power=140, Cycle/Burst=200, Time=240 seconds. Chromatin extracts were immunoprecipitated with Gata1 antibodies overnight at 4°C, and then incubated with protein A Dynabeads (Invitrogen) for another 4 hours. ChIP complexes were eluted and cross-linking reversed by heating at 65°C overnight. DNA was extracted with DNA Clean and Concentrator columns (Zymo Research).

For library preparation, Biotin labeled post-ChIP DNA was captured by 5 μl Streptavidin C-1 beads (Thermo Fisher), and followed by Tn5 treatment. The amount of Tn5 used and the number of PCR cycles performed was based on the post-ChIP Qubit amounts, as previously described (Mumbach et al., 2016). HiChIP libraries were size-selected for 300-800bp fragments. All libraries were sequenced on the Illumina Nextseq 500 to an average depth of 200 million total reads.

#### RNA-seq data analysis

RNA-seq reads were aligned to the reference mouse genome mm10 using STAR (Dobin et al., 2013) with default parameters. Aligned reads were counted in the genomic transcripts annotations from GenomicFeatures (Lawrence et al., 2013) using Rsamtools (Morgan M, 2016). The differentially expressed gene analysis was performed by using DESeq2 (Love et al., 2014), with the threshold at adjusted *P*-value 0.01 and fold-change 2. Benjamini-Hochberg multiple hypothesis testing correction was performed.

#### ATAC-seq data analysis

ATAC-seq reads were trimmed for adapter sequences and low alignment quality using Cutadapt (Martin, 2011), with the parameters -q 30 --minimum-length 36. Paired-end reads were aligned to the mm10 reference genome by using Bowtie2 (Langmead et al., 2009), with the parameter -X 2000 allowing fragments of up to 2kb to be aligned (Buenrostro et al., 2015). ATAC-seq peaks were called using MACS2 (Zhang et al., 2008) with following parameters (--nomodel --nolambda --keep-dup all --call-summits) (Buenrostro et al., 2015). Peak calling was done in two biological replicates respectively with the same parameter setting. One replicate of each cell type was used for subsequent analysis. We then extended the summits to 500bp window (±250 bp) as peaks and filtered using the consensus excludable ENCODE blacklist (Consortium, 2012).

#### ChIP-seq data analysis

ChIP-seq reads were aligned to the mm10 reference genome by using Bowtie2 (Langmead et al., 2009) with default parameters. Duplicate reads were removed using PICARD tools (http://picard.sourceforge.net). ChIP-seq peaks were called by using MACS2 (Zhang et al., 2008) with following parameters (--nomodel --keep-dup 1 --extsize=146 -q 0.01). Peaks were filtered using the consensus excludable ENCODE blacklist (Consortium, 2012). The heatmap or density plot for ChIP-seq signal were plotted based on the binned density matrix range from ±2.5kb centered by the summit generated by using the CEAS software (Shin et al., 2009). Peak calling was done in two biological replicates with the same parameter setting. The replicates were merged if they are available and shown with high correlation. The datasets of Gata1, H3K27ac, p300 and Myb ChIP-seq in MEL cells were downloaded from ENCODE (Consortium, 2012).

#### HiChIP data analysis

HiChIP datasets were processed using HiC-Pro (Mumbach et al., 2016; Mumbach et al., 2017; Servant et al., 2015) with default settings. Specifically, HiChIP paired-end reads were aligned using Bowtie2 with --very-sensitive -L 30 --score-min L, −0.6, −0.2 --end-to-end --reorder. The restriction sites were obtained by scanning MboI restriction fragments across mouse genome (mm10). Specific configuration file (config-hicpro.txt) can be found at https://bitbucket.org/qzhu/wenqing-hichip/src/default/hicpro/config-hicpro.txt. Valid interactions pairs (named valid Pairs) were converted to a .hic file using hicpro2juicebox.sh script (utility tool of HiC-Pro). The generated .hic file contains interaction matrices at fragment and base pair resolutions. Interaction maps were visualized with Juicebox (Durand et al., 2016a).

#### HiCHIP loop calling

The HiCCUPS function of Juicer-tools 1.7.6 was used to call high-confidence loops based on a .hic file (Rao et al., 2014). We set -m 512 -r 5000,10000,25000 -f 0.2,0.2,0.2 -p 4,2,1 -k VC_SQRT -i 7,5,3 -d 20000,20000,50000 --ignore_sparsity -c 1,2,3,4,5,6,7,8,9,10,11,12,13,14,15,16,17,18,19,X,Y -t 0.02,1.5,1.75,2. This combination of settings finds total loops at the 5-kb, 10-kb, and 25-kb resolutions using the interaction matrices at matched resolutions, and also applied the appropriate normalization step for interaction matrix (vanilla coverage square-root, VC SQRT). Loops at these resolutions were subsequently merged.

#### Definition of proximal and distal regions

Proximal regions or promoters are ±2kb windows centered by RefSeq transcription start sites (TSS) locations (O’Leary et al., 2016), whereas the remaining regions of the genome were considered as distal regions.

#### Identification of differential peaks

The differential peaks (from ChIP-seq or ATAC-seq) between EryP and EryD were determined as previously described (Huang et al., 2016; Xu et al., 2012), using MAnorm (Shao et al., 2012), with the threshold of M-value 1 and FDR 0.01. We termed differential peaks as EryP-/EryD-specific peaks, whereas the remainders were considered as shared peaks.

#### Motif enrichment analysis

To identify enriched motifs within a set of enhancers, we performed motif enrichment analysis using HOMER (Heinz et al., 2010), as previously described (Huang et al., 2016). The position weight matrixes (PWM) of core vertebrate motifs were downloaded from the JASPAR database (Mathelier et al., 2014). The enrichment score of each motif in a set of enhancer regions was defined as −log_10_(*P*-value), where *P*-value corresponds to the significance of observed over-representation of each motif site in enhancer regions compared to control regions randomly selected from the genome, which is calculated based on hypergeometric distribution.

#### Enhancer-promoter loops comparison

The total number of enhancer-promoter loops (E-P loops) between EryP and EryD conditions were comapred at each 10-kb, and 25-kb resolution, respectively. The enhancers were defined from GATA1 ChIP-seq peaks, while the promoter was defined as TSS±2kb region. Before comparison, neighboring GATA1 ChIP-seq peaks were merged, using BEDOPS (Neph et al., 2012a), to keep the resolution of ChIP-seq data consistent with the resolution of HiChIP data. For example, for 10-kb loop comparison, we merged ChIP-seq peaks at a threshold of 10-kb. The number of enhancer-promoter loops was then tallied in each EryP and EryD cells, for EryP-specific genes, EryD-specific genes, and common genes. Common genes were defined as non EryP-/EryD-specific genes and with relatively expression in both EryP and EryD cells based on RNA-seq distribution. The difference in the total number of enhancer-promoter loops across conditions was computed. To test its significance, we picked 1,000 genes randomly in the genome to compute the difference in the number of enhancer-promoter loops across EryP and EryD conditions. 1,000 permutations of random genes were analyzed to derive an empirical *P*-value for the difference (EryP-EryD).

#### Virtual 4C visualization

Virtual 4C analysis is used to examine the number of interactions formed with an anchor region of interest (Mumbach et al., 2017). Based on the defined anchor in our case (the E1 enhancer in *Hemgn* locus and E2 enhancer in *Tnfaip2* locus), we computed the number of interactions formed with the anchor site using the Juicer-tools dump function (Durand et al., 2016b). The specific settings were “observed <hicfile> <range1> <range2> FRAG 1” which means observed number of reads at 1-fragment resolution, range1 specifies the anchor site, range2 specifies ±400 frag from the anchor site. Afterward, reads at fragment resolution were converted to basepair resolution by checking back the restriction-site file. Reads were redistributed across 200 basepair bins at a bin size of 750bp to give the final virtual 4C plot. Note that fragment resolution dump was required as the minimal resolution of basepair interaction matrix in a hic file was 5-kb, while at fragment resolution 1 fragment is equivalent to about 400bp.

#### Average profiles of enhancer-promoter interactions

A summary virtual 4C plot was generated to summarize the enhancer-promoter interactions for over EryP- and EryD-specific genes in EryP and EryD cells, respectively. We treated each promoter as anchor site in each case, where a promoter is defined as a H3K4me3 peak at TSS±2kb. For enhancer definition, we used GATA1 ChIP-seq peaks. Next, for each enhancer neighboring the promoter at a distance of ±100kb, we tabulated the number of interaction of reads that each enhancer forms with the promoter in the HiChIP dataset. Juicer-tools dump function (Durand et al., 2016b) was used to provide the number of reads at fragment resolution, and then were converted to base pair resolution. Reads were summarized to the corresponding enhancer bin based on the ordinal position of enhancer relative to promoter (1^st^ bin if reads are for first enhancer from promoter, 2^nd^ bin if reads are for 2^nd^ enhancer from promoter, and so on). Average reads per enhancer bin were computed, giving **Fig. 4D**.

#### Data visualization

The ChIP-seq and ATAC-seq signal tracks were visualized using Integrative Genomics Viewer (IGV) (Robinson et al., 2011). HiChIP interaction maps were visualized with Juicebox (Durand et al., 2016a), and HiChIP loops were visualized with WashU genome browser (Zhou et al., 2011).

#### Processing of ENCODE mouse datasets

H3K27ac ChIP-seq and RNA-seq datasets of mouse tissues during development were downloaded from ENCODE (Consortium, 2012). To balance the various sequencing depth among samples, H3K27ac and RNA-seq raw data were sampled to 10 and 30 million reads, respectively. All tissue samples, in which the H3K27ac ChIP-seq and RNA-seq data were both available in each instance, were processed in an unbiased way, and were further filtered through the following criteria: 1) the sequencing depth at least more than 10 million per sample; 2) at least 3 developmental time points per tissue; 3) with at least 2 biological replicates available. 42 embryonic samples, including heart, liver, forebrain, midbrain, hind brain, limb, and embryonic facial prominence, and covering different developmental stages from E10.5 to E16.5, successfully went through all filtrations. For each tissue, we grouped the available samples into three categories labeled as “Early”, “Middle”, and “Late” based on their annotated developmental ages (**Fig. 6B**). In brief, the beginning 1-2 developmental ages and the last 1-2 developmental ages were grouped into “Early” and “Late” respectively, and remaining were considered as “Middle” (**Fig. 6B**).

We construct the gene expression matrix in ENCODE 42 samples, using log_2_(RPKM+1) from RNA-seq data. For each cell type, cell-type-specific expressed genes were defined as the top 500 genes, ranked based on the cell-type-specificity of gene expression. We constructed the enhancer profiles matrix in three steps. 1) We defined distal H3K27ac peaks as enhancers for each sample. 2) An enhancer catalog across all cell types was curated based on the union of the enhancers in each cell type, followed by a merge of overlapping enhancers to remove redundant ones. 3) H3K27ac density (log_2_(RPKM+1)) of enhancer catalog in each cell type was used to represent the enhancer profiles matrix. For each cell type, cell-type-specific enhancers were defined as the top 5,000 enhancers ranked by their cell-type-specificity, which was calculated as the fold-enrichment of H3K27ac density compared with the average H3K27ac density across all cell types. The gene expression matrix and enhancer profile matrix of ENCODE 42 samples were used for C-score analysis described below.

#### Processing of Roadmap human datasets

The H3K27ac ChIP-seq and RNA-seq data in 48 human cell types were downloaded from Roadmap (Bernstein et al., 2010). We manually classified 48 cell types into three groups “Embryonic”, “Fetal”, and “Adult” based on their annotation information. As with the processing of ENCODE datasets (Consortium, 2012), we constructed gene expression matrix and enhancer profile matrix, and then defined cell-type-specific enhancers and genes in Roadmap 48 cell types for C-score analysis described below (**Fig. 6A**).

#### C-score analysis

We introduced a computational metric, C-score, to systematically assess the association between a set of genomic regions and a set of cell-type-specific expressed genes. This stratey was adopted from GREAT analysis, which predicts functions of *cis-*regulatory regions by analyzing the annotations of the nearby genes (McLean et al., 2010). As illustrated in **Fig. S2A**, two steps, “assignment” and “assessment”, were involved in C-score calculation. In “assignment” step, a set of genomic regions, *e.g.* ATAC-seq or ChIP-seq peaks, were mapped to candidate genes within ±50kb using the ‘nearest neighbor gene’ approach (Huang et al., 2016; Xu et al., 2012) using HOMER (Heinz et al., 2010) (**Fig. S2A**). In “assessment” step, we performed Fisher’s exact test to assess the enrichment significance of a set of genomic region assigned cell-type-specific genes, where −log_10_*(P*-value) was termed as the C-score. C-scores ≥ 200 were presented as 200 to create appropriate color range in a heatmap. In EryP/EryD dataset, we used the same number of cell-type-specific genes (943 genes) in C-score analysis to exclude potential biases on *P*-value due to the size of the gene set. In ENCODE and Roadmap datasets, top 5,000 cell-type-specific enhancers and top 500 cell-type-specific genes were used in C-score analysis. C-score analyses were performed in replicates independently and the results were highly consistent.

#### Validation of C-score analysis

Parameters in “assignment” and “assessment” steps (**Fig. S2A**) may bia C-score analysis and these parameters are 1) various mapping distance (±25kb or ±100kb), 2) multiple genes mapping and 3) scoring metric (**Fig. S2A**, **middle**). To exclude false positives, we validate C-score analysis by repeating the analysis with one parameter changed at a time (**Fig. S2A**, **right**). Within “different mapping distances”, the nearest genes within ±25kb or ±100kb of enhancer center were designated as the enhancer target genes; In “multiple-genes mapping” approach, all genes within ±50kb of enhancer center were designated as the enhancer target genes. To validate the association results scored by Fisher’s exact test, we applied the Binomial test, which was used in GREAT analysis (McLean et al., 2010), as an alternative scoring metric, and termed this score as GREAT score (G-score) (**Fig. S2A**). To calculate a G-score, we annotated an genomic region (±50kb from TSS) to a EryP- or EryD-specific gene (McLean et al., 2010). Then, the enrichment significance of a set of EryP-/EryD-specific peaks (ATAC-seq or ChIP-seq peaks) across annotated genomic regions was assessed by using Binomial test, where −log_10_(*P*-value) was defined as the G-score (**Fig. S2A right**). As with C-score, G-scores increased as the association of genomic regions to specific expressed genes becomes greater. We performed the validation of C-score analysis on all datasets, and achieved consistent C-score patterns as compared with the initial C-score analysis. Representative analyses with EryP-/EryD-specific active enhancers, EryP-/EryD-specific Gata1 peaks and with Roadmap datasets were summarized in **Fig. S2.**

#### Enrichment analysis of GWAS SNPs

The SNPs curated in GWAS Catalog (Welter et al., 2014) were downloaded through the UCSC Table Browser (Karolchik et al., 2004). In 48 human cell types (Roadmap), we computed enrichment of SNPs on cell-type specific enhancers (**Fig. 6A**) and on C-score-related enhancers, in which enhancers assigned genes were overlapped with cell-type specific genes (**Fig. 6A** and **6E**). In Brief, for each group of enhancers, the enrichment of SNPs was defined as the fold enrichment relative to genomic background. It was calculated as following: (m/n)/(M/N), where m and M represent the number of within-group and genome-wide SNPs respectively, and n and N represent the number of within-group and genome-wide loci respectively. The fraction of enhancers with SNPs represents the percentage of enhancer that overlaps with at least 1 SNP.

#### Lentiviral shRNA infection

10^6^ MEL cells were mixed with virus particles generated by Dox-inducible *Myb shRNAs*, and control vector (Sigma), supplemented with 10 μg/ml polybrene (Sigma-Aldrich). Lentiviral vectors bearing the coding sequence of the Puromycin-resistance gene permitted a positive selection of 2 μg/ml Puromycin (Sigma-Aldrich) treatment. shRNA sequences are listed in supplemental **Table S3**.

#### CRISPR/Cas9 enhancer deletion in MEL cells and mESCs

To generate biallelic deletion of Enhancer 1 (E1, chr4: 46410632-46411247, in *Trmo*, *Hemgn*, *Anp32b* and *Nans* locus), sgRNAs targeting 5’- and 3’-flanking regions of E1 were designed and synthesized respectively. sgRNA sequences are listed in supplemental **Table S4**. Two overlapping oligonucleotides carrying sgRNA sequence targeting 5’-flanking region and two overlapping oligonucleotides carrying sgRNA targeting 3’-flanking region were annealed and cloned respectively, as previously described (Bauer et al., 2015). In brief, 10 μM guide sequence oligos and 10 μM complement oligo were mixed with 1X ligation buffer supplemented with 5 U of T4 Polynucleotide Kinase (PNK) in 10 μl reaction. Anneal in a thermocycler using 37°C for 30 min; 95°C for 5 min and then ramp down to 25°C at 0.1°C/sec. The annealed oligos were then ligated into pX330 vector using a Golden Gate assembly. Ligation mixture [100 ng vector, 1 μM annealed oligos, 40 U BbsI restriction enzyme (NEB), 1 mM ATP, 0.1 mg/ml BSA and 750 U T4 DNA ligase (NEB), and 1X restriction enzyme buffer] were incubated in a thermocycler using 20 cycles of 37°C for 5 min, 20°C for 5 min; followed by 80°C for 20 min.

pX330 construct with sgRNA targeting 5’-flanking region and px330 construct with sgRNA targeting 3’-flanking region were co-transfected with pCas9-GFP (Addgene #44719) at the ratio of 1:1:2 into MEL cells or mESCs by Lipofectamine 2000 (Invitrogen). The top 5% of GFP^+^ cells was isolated 48 hours post-transfection by FACS. Single cell derived colonies were screened for biallelic deletion of targeting region. For mESCs, FACS isolated GFP^+^ cells were plated on feeder cells to generate single cell colonies. Genotyping primer sequences are listed in supplemental **Table S5**.

Biallelic deletion of Enhancer 2 (E2, chr12: 111517811-111518523, in *Tnfaip2* and *Eif5* locus) in MEL cells and mESCs were generated using a similar strategy with sgRNA**s** targeting flanking regions of E2. sgRNA sequences are listed in supplemental **Table S4**. Genotyping primer sequences are listed in supplemental **Table S5**.

#### RNA extraction and qRT-PCR

Total RNA was extracted with TRIzol (Thermo Fisher) and reverse transcribed to cDNA with QuantiTect Reverse Transcription Kit (Qiagen) according to the manufacturer’s instructions. cDNA samples were subjected to qRT-PCR using the iQ SYBR Green Supermix (Bio-Rad) in the CFX384 Touch Real-Time PCR Detection System(Bio-Rad). Primer sequences are listed in supplemental **Table S6**. Values are expressed as log_10_2^DeltaCt using *Actin beta (Actb)* as a control gene.

#### Quantification and Statistical Analysis

Each experiment was repeated at least twice using a minimum of three biological replicates per condition. Statistical analysis was performed with unpaired Student’s *t-*test. Error bars indicate the S.E.M.; n=3 in **Fig. 4, Fig. S1, Fig. S4** and **Fig. S5**. In box plots in **Fig. 6** and **Fig. S2**, centerline of each box represents the median, the box limits represent the 25th and 75th percentiles and the whiskers represent the 5th and 95th percentiles. Statistical analysis of **Fig. 4E, Fig. S5C** and **S5D** were performed with permutation test in 1,000 random genes. *P*-values were calculated and statistical significance indicates \**P*<0.05, \*\**P*<0.01, \*\*\**P*<0.001.

#### Data Availability

The RNA-seq, ChIP-seq, ATAC-seq, and HiChIP data generated in this study have been deposited in Gene Expression Omnibus (GEO) under accession number GSE112717.

