## Supplementary material for "Enhancer-dependence of gene expression increases with developmental age": All supplemental information

**<u>Contents:</u>**


- **Supplemental Figures and Legends (Fig. S1-S6)**
- **Supplemental Table S1**
- **Supplemental Table S2**
- **Supplemental Table S3**
- **Supplemental Table S4**
- **Supplemental Table S5**
- **Supplemental Table S6**

### Supplemental Figures and Legends

#### Figure S1, related to Fig. 1

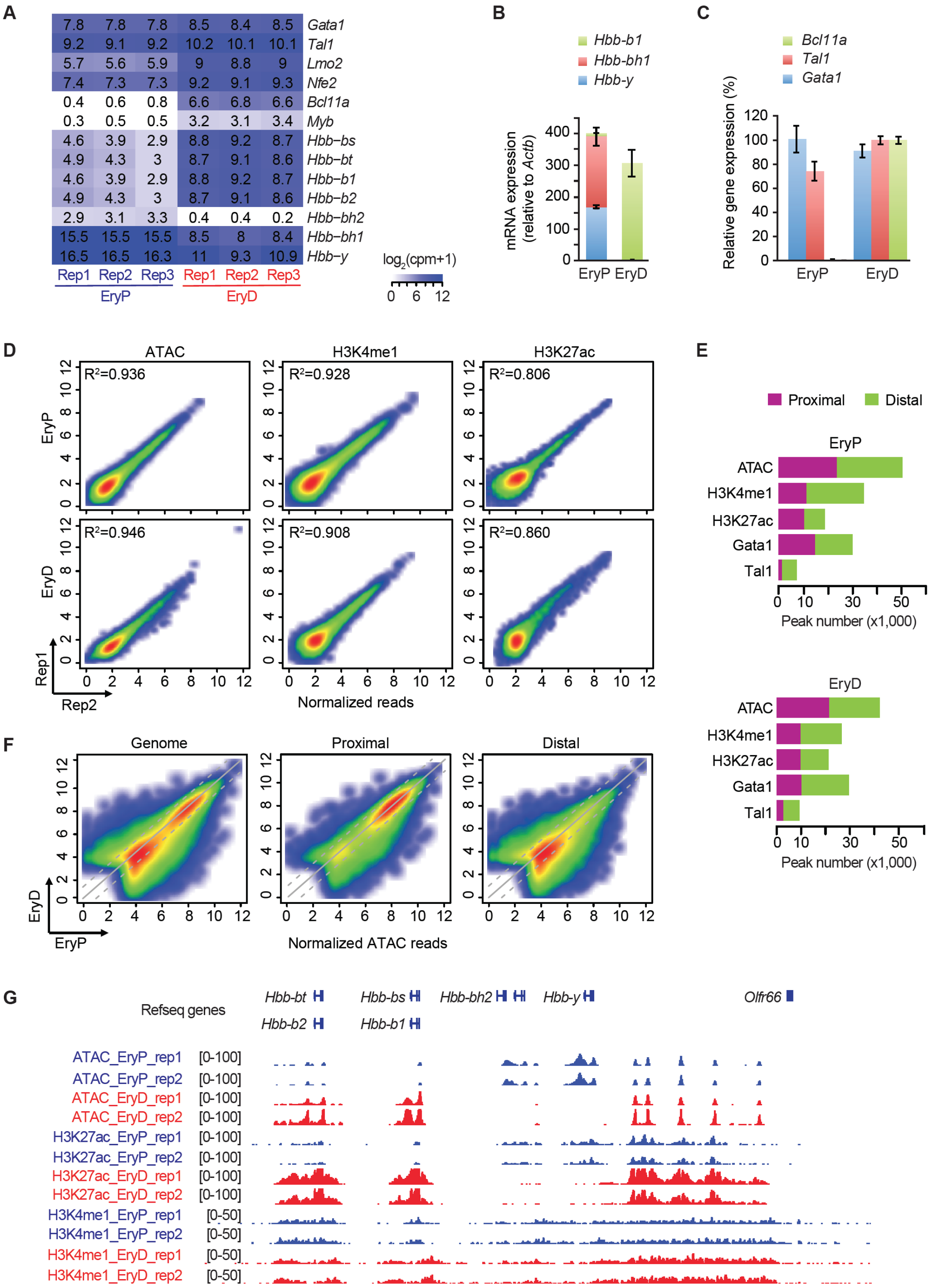

**Figure S1, related to Fig. 1: Genome-wide comparative analysis of EryP and EryD**

**(A)** Heatmap of RNA-seq analysis of selected erythroid specific genes in EryP and EryD.

**(B)** qRT-PCR analysis of *Hbb-b1*, *Hbb-bh1*, and *Hbb-y* in EryP and EryD.

**(C)** qRT-PCR analysis of *Gata1*, *Tal1*, and *Bcl11a* in EryP and EryD.

**(D)** Reproducibility analysis of ATAC-seq and ChIP-seq of histone marks. Density plots of normalized ATAC-seq or ChIP-seq reads from two biological replicates show reproducibility of ATAC-seq and ChIP-seq data. *x*-axis and *y*-axis represent ATAC-seq or ChIP-seq normalized reads ($\log_2$(RPKM+1)) at promoters (±2kb region from a transcription start site (TSS)).

**(E)** The genomic distribution and total peak numbers of ATAC-seq and ChIP-seq of histone marks and transcription factors in EryP and EryD, respectively.

**(F)** Density plots show the comparison of ATAC-seq signal ($\log_2$(RPKM+1)) in EryP and EryD at genome-wide (left), proximal (middle) and distal (right) regions. ATAC-seq signals are normalized reads within the ATAC peaks detected in EryP or EryD.

**(G)** Representative IGV snapshot of chromatin landscapes in LCR region.

Experiments were replicated at least twice in **B** and **C**. **P*<0.05, ***P*<0.01, ****P*<0.001, unpaired one-tailed Student's *t*-test. Error bars indicate the S.E.M.; n=3 (**B** and **C**).

Figure S2, related to Fig. 1, 2, 3 and 6

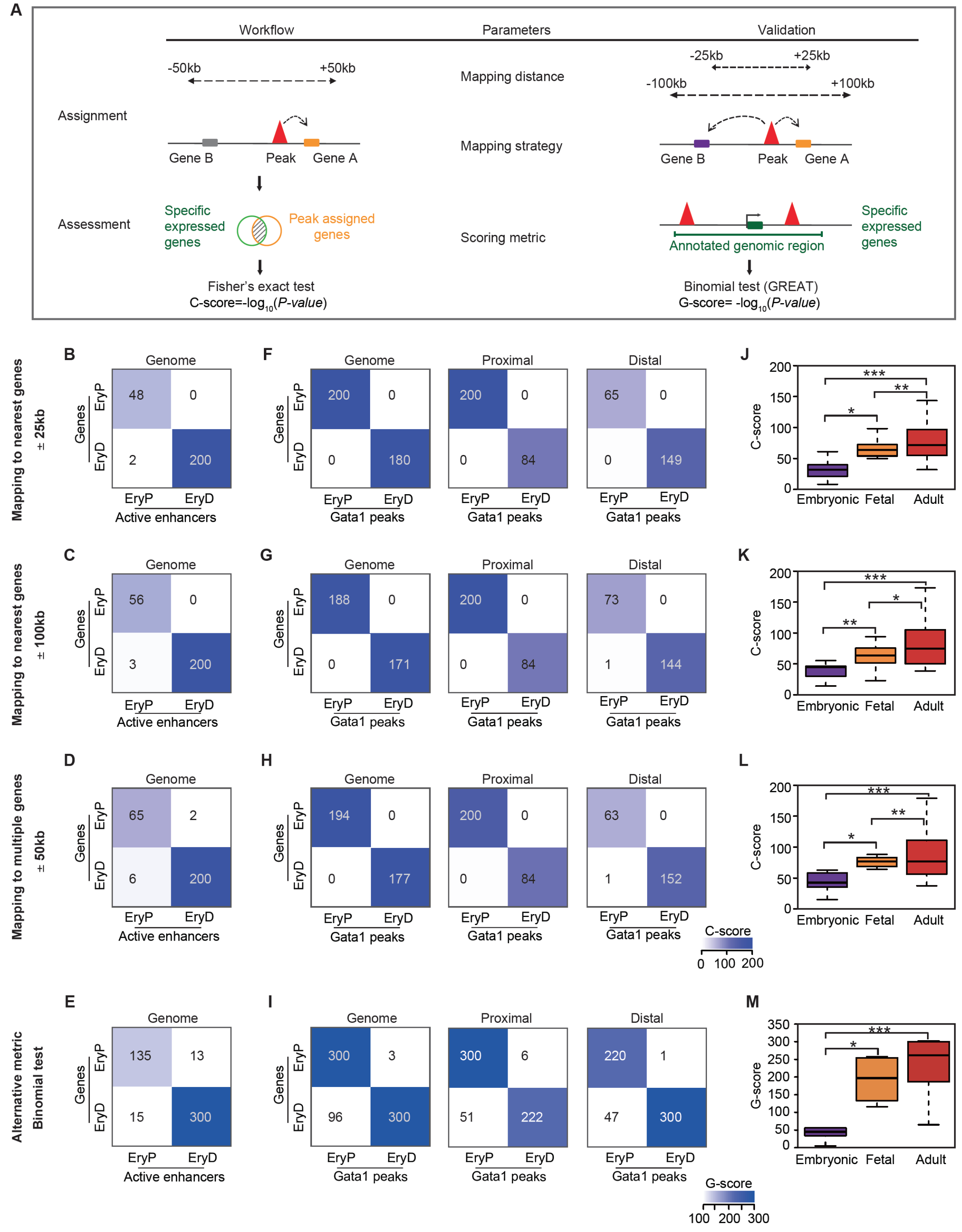


### Figure S2, related to Fig. 1, 2, 3 and 6: Definition and validation of C-score

**(A)** Diagram of the definition and validation of C-score.

**(B-D)** Validation of C-score analyses using EryP-/EryD-specific active enhancers and gene expression. Validation of “mapping distance” is in **B** (±25kb) and **C** (±100kb). Validation of “mapping to multiple genes” is in **D**. Rows are EryP-/EryD-specific genes. Columns are EryP-/EryD-specific active enhancers (also see **Fig. 2E**, **Methods**).

**(E)** Association analysis of EryP-/EryD-specific active enhancers and gene expression using an alternative scoring metric, Great-score (G-score). A G-score, calculated by using Binominal test, was designated to assesse the enrichment significance of a set of EryP-/EryD-specific enhancers (or cell-type-specific ChIP-seq peaks) across the annotated genomic regions (highlighted in green) (**Fig. S2A right, Methods**). Rows are EryP-/EryD-specific genes. Columns are EryP-/EryD-specific active enhancers. Also see **Fig. 2E**.

**(F-I)** Validation of C-score analyses using EryP-/EryD-specific Gata1 peaks and gene expression at genome-wide, proximal and distal regions. Validation of “mapping distance” is in **F** and **G**. Validation of “mapping to multiple genes” is in **H**. Validation using Binomial test is in **I**. Also see **Fig. 3D, 3E**.

**(J-M)** Validation of C-score analyses using cell-type-specific enhancers to gene expression in Roadmap 48 human cell types. Validation of “mapping distance” is in **J** and **K**. Validation of “mapping to multiple genes” is in **L**. Validation using Binomial test is in **M**. C-Score or G-score of individual cell type are summarized within each classification (also see **Fig. 6F**, **Methods**).

In box plot, the box represents the 25th and 75th percentiles, while the whiskers represent the 5th and 95th percentiles. *$P$<0.05, **$P$<0.01, ***$P$<0.001, unpaired one-tailed Student’s *t*-test (**J, K, L** and **M**).

Figure S3, related to Fig. 2

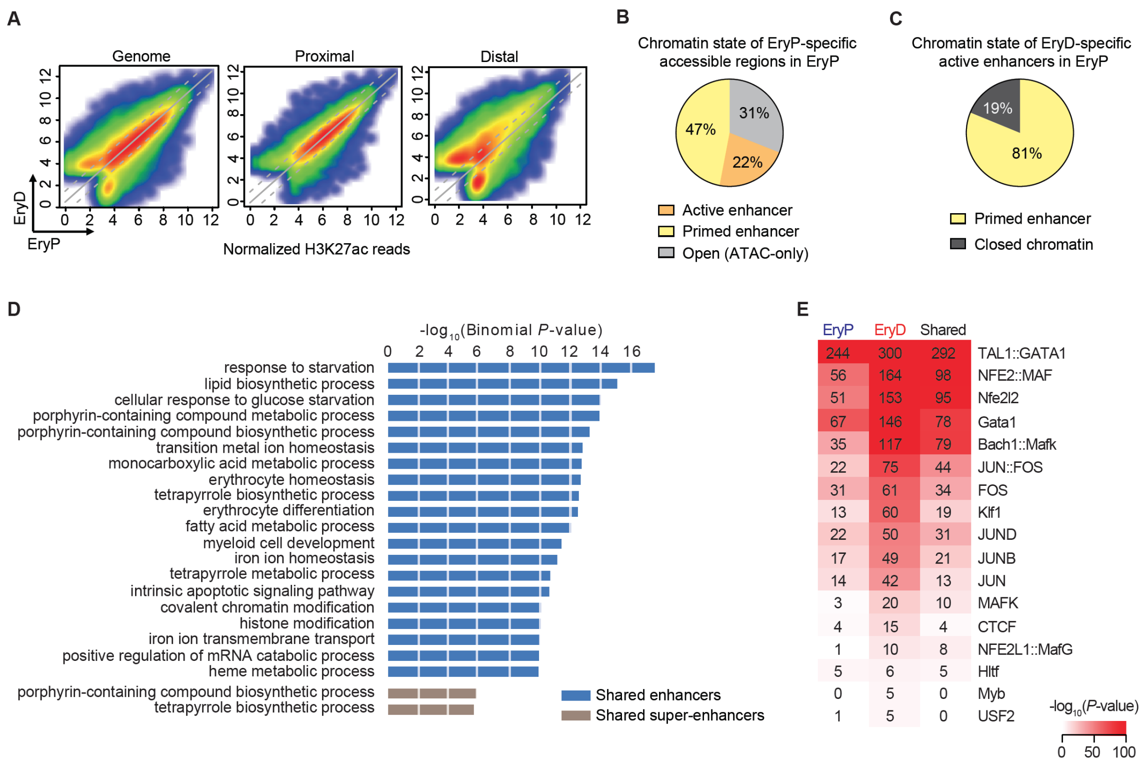


**Figure S3, related to Fig. 2: Chromatin state of EryP-/EryD-specific distal accessible regions.**

**(A)** Density plots show the comparison of the H3K27ac ChIP-seq signal in EryP and EryD at genome-wise (left), proximal (middle) and distal (right) regions. H3K27ac ChIP-seq signals are normalized reads within the H3K27ac peaks detected in EryP or EryD.

**(B)** Chromatin state of EryP-specific distal accessible regions in EryP cells. Pie charts show the percentage of “primed enhancer”, “active enhancer” and “ATAC-only” of each group, based on the presence of H3K27ac peaks or H3K4me1 peaks. Primed enhancer is marked with H3K4me1, lacking H3K27ac, whereas active enhancer is marked with both H3K4me1 and H3K27ac.

**(C)** Chromatin state of EryD-specific active enhancer in EryP cells. Pie charts show the percentage of “primed enhancer” and “closed chromatin” of each group, based on the presence of ATAC peaks, H3K27ac peaks or H3K4me1 peaks. Primed enhancer is marked with H3K4me1, lacking H3K27ac, whereas closed chromatin is unmarked with ATAC peaks.

**(D)** GREAT functional enrichment analysis of EryP- and EryD- shared enhancers and shared super-enhancers.

**(E)** Motif enrichment analysis of EryP-, EryD-specific and shared active enhancers. Gata1 and Tal1 motif were enriched in both EryP and EryD active enhancers.

Figure S4, related to Fig. 3

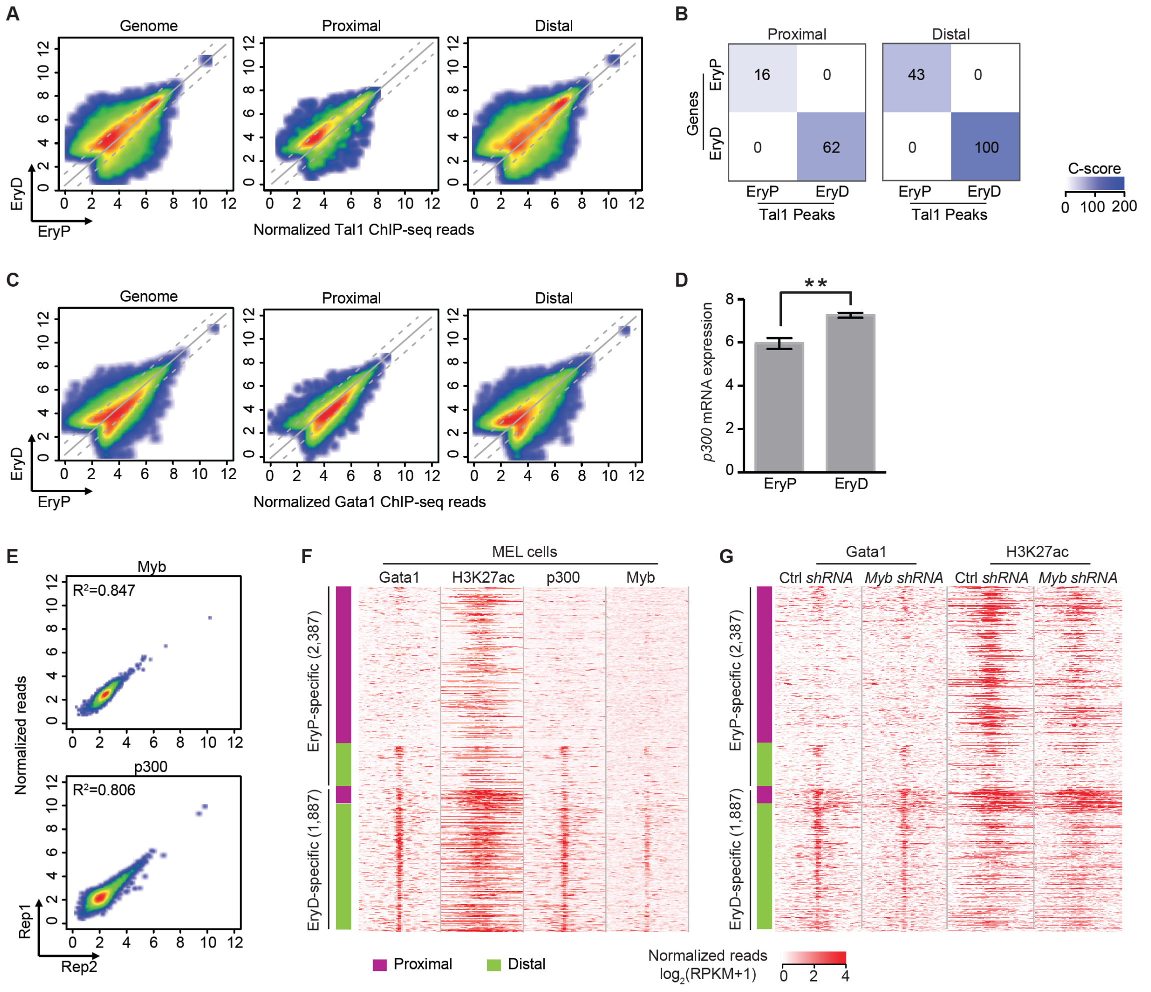


**Figure S4, related to Fig. 3: Genomic occupancy of master transcription factors in EryP and EryD**

**(A-B)** Density plots show the comparison of Tal1 ChIP-seq signal in EryP and EryD at genome-wide (left), proximal (middle) and distal (right) regions. Tal1 ChIP-seq signals are normalized reads within the Tal1 peaks detected in EryP or EryD. Corresponding association studies of EryP-/EryD-specific Tal1 peaks and specific genes at proximal (left) and distal (right) regions are shown in (**B**).

**(C)** Density plots show genome-wide comparison of Gata1 ChIP-seq signal in EryP and EryD at genome-wide (left), proximal (middle) and distal (right) regions. Gata1 ChIP-seq signals are normalized reads within the Gata1 peaks detected in EryP or EryD.

**(D)** RNA-seq analysis of *p300* in EryP and EryD.

**(E)** Density plots of normalized ChIP-seq read from two biological replicates show reproducibility of Myb and p300 ChIP-seq data. *x*-axis and *y*-axis represent ChIP-seq normalized reads ($\log_2$(RPKM+1)) at promoters (±2kb region of a transcription start site (TSS)) from two biological replicates.

**(F)** Heatmap of the ChIP-seq normalized read density of Gata1, H3K27ac, p300 and Myb in MEL cells around the Gata1 peak summit in EryP and EryD, shown in **Fig. 3G.**

**(G)** Heatmap of the normalized ChIP-seq read density of Gata1 and H3K27ac in MEL cells treated with control *shRNA* (Ctrl *shRNA*) or *Myb shRNA,* around the Gata1 peak centers in EryP and EryD, shown in **Fig. 3G**.

**P*<0.05, ***P*<0.01, ****P*<0.001, unpaired one-tailed Student's *t*-test. Error bars indicate the S.E.M.; n=3 (**D**).

**Figure S5, related to Fig. 4**

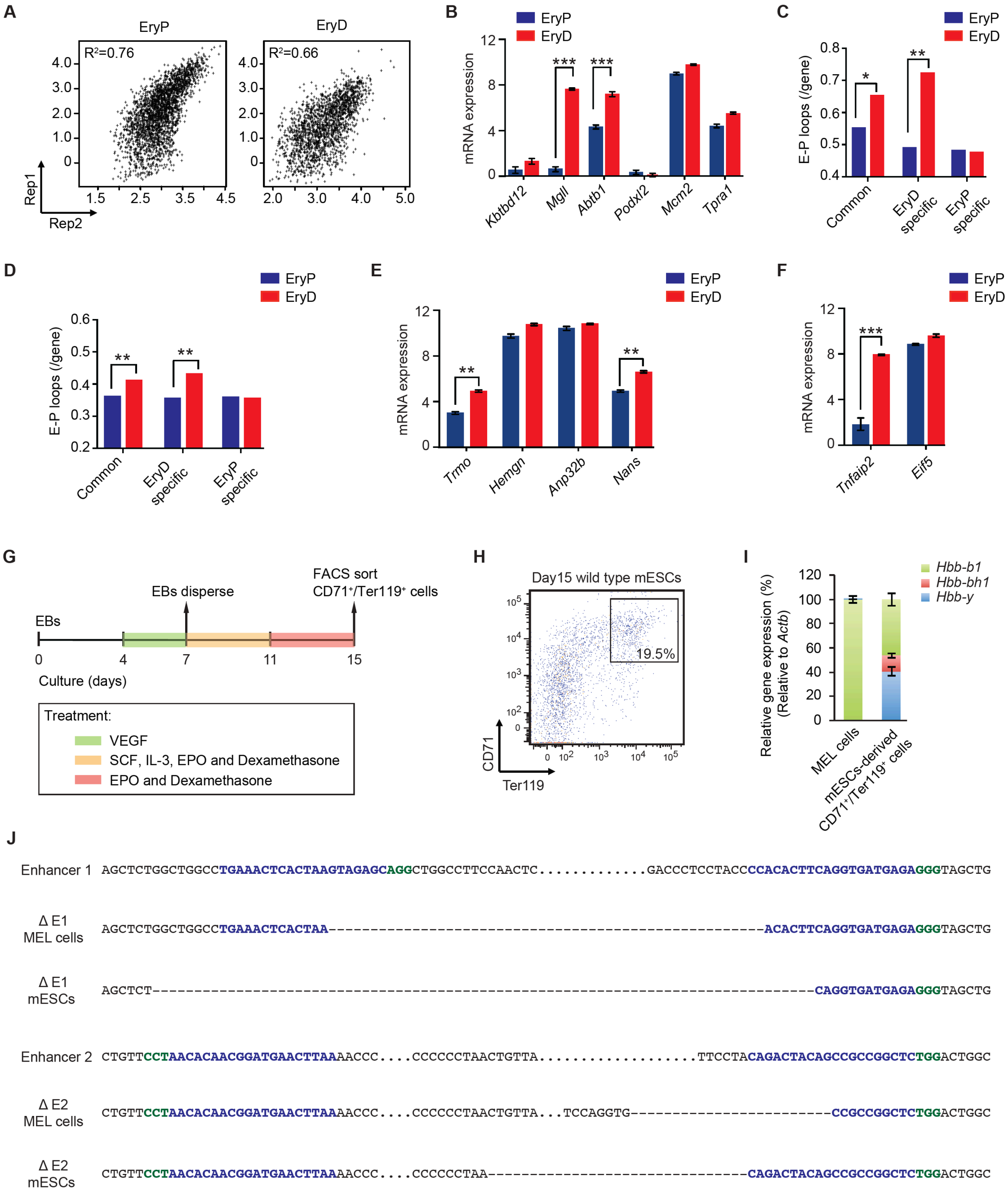

**Figure S5, related to Fig. 4: Gata1 HiChIP and enhancer deletion in primitive and definitive cells.**

**(A)** Gata1 HiChIP reproducibility scatter plots. Reads reproducibility of Gata1 HiChIP from two biological replicates is shown in EryP cells (left) and EryD cells (right), respectively.

**(B)** RNA-seq analysis of *Kbtbd12, Mgll, Abtb1, Podxl2, Mcm2* and *Tpra1* in EryP and EryD.

**(C-D)** Quantification of enhancer-promoter (E-P) loops in EryP and EryD cells. Comparisons were in three groups, EryP-specific genes, EryD-specific genes, and common expressed genes. Comparison between rep-2 of EryP and EryD at 25-kb resolution is in (**C**). Comparison between rep-1 of EryP and EryD at 10-kb resolution is shown in (**D**).

**(E)** RNA-seq analysis of *Trmo, Hemgn, Anp32b* and *Nans* in EryP and EryD.

**(F)** RNA-seq analysis of *Tnfaip2* and *Eif5* in EryP and EryD

**(G)** Schematic of experimental timeline of mESCs differentiation.

**(H)** Representative flow cytometry profile of CD71 and Ter119 in day 15 wild-type mESCs.

**(I)** qRT-PCR analysis of *Hbb-b1*, *Hbb-bh1*, and *Hbb-y* in MEL cells and mESCs-derived $CD71^+$/$Ter119^+$ cells. Day 5 differentiated MEL cells and FACS isolated $CD71^+$/$Ter119^+$ cells from day 15 mESCs were collected for analysis.

**(J)** Genotypes of ΔE1 and ΔE2 in MEL cells and mESCs generated by CRISPR/Cas9 editing. Sanger sequencing confirmed the presence of genome editing at the targeted site (blue bold letters), which resulted in biallelic enhancer deletion. Dash line represents deleted nucleotides. PAM sequence is green and bold.

Experiments were replicated at least twice in **H** and **I**. Error bars indicate the S.E.M.; n=3 (**B**, **E**, **F** and **I**). *$P<0.05$, **$P<0.01$, ***$P<0.001$, unpaired one-tailed Student's *t*-test (**B, E** and **F**). *P*-value in **D** and **E** represents permutation test in 1,000 random gene selection of matched size.

**Figure S6, related to Fig. 6**

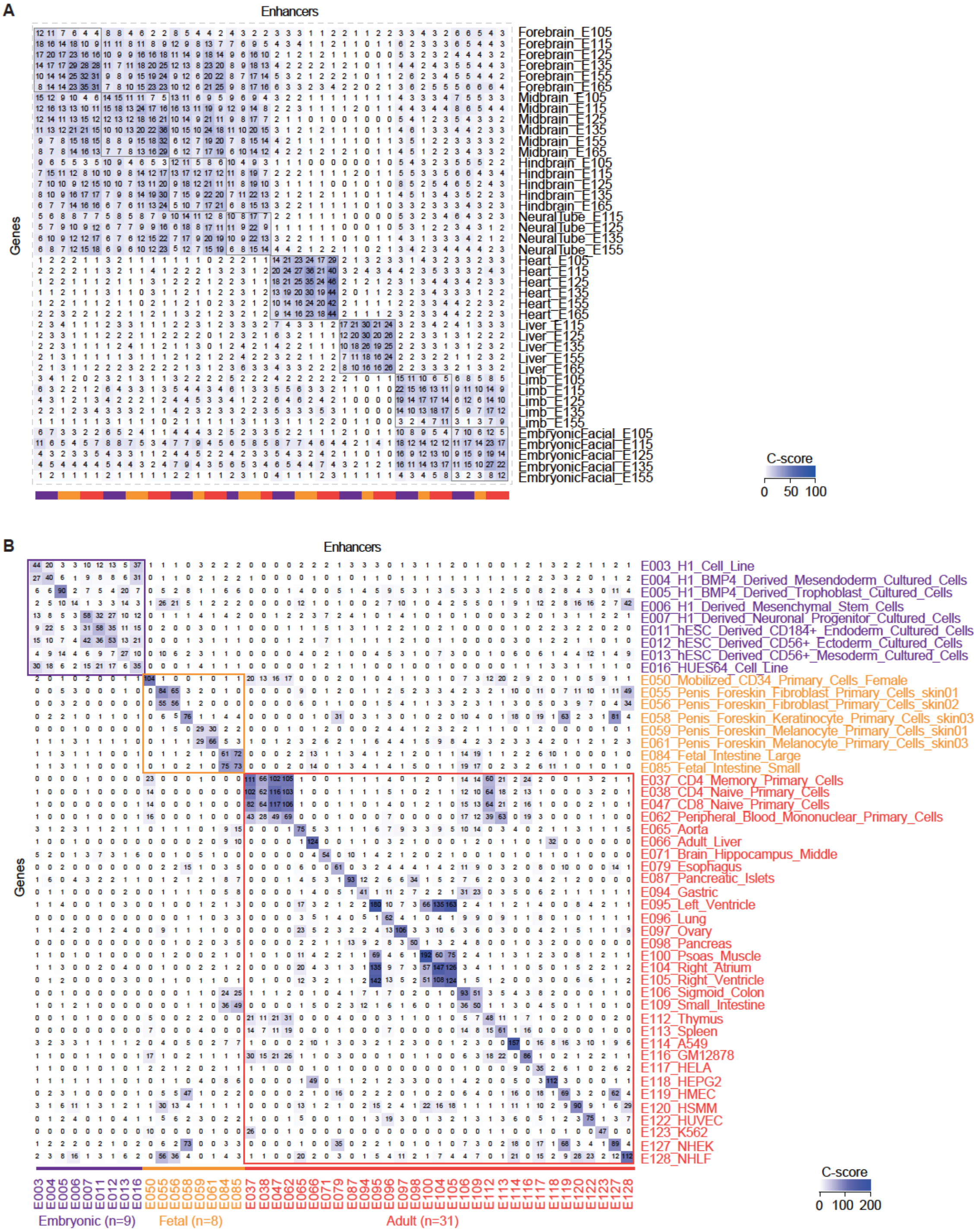

**Figure S6, related to Fig. 6: Association studies of cell-type-specific enhancers and gene expression during mouse and human ontogeny**

**(A)** The C-score matrix from association studies in mouse ENCODE datasets. This panel is similar to **Fig. 6C**, but with detailed annotations.

**(B)** The C-score matrix from association studies in human Roadmap datasets. This panel is similar to **Fig. 6E**, but with detailed annotations.

**Supplemental Table S1: RNA-seq, ATAC-seq, ChIP-seq and HiChIP profiling datasets generated in this study**

| Dataset | Data type | Cell type | GEO ID |
|---|---|---|---|
| RNAseq-EryP-Rep1 | RNA-seq | Primitive erythroblast | GSM3081986 |
| RNAseq-EryP-Rep2 | RNA-seq | Primitive erythroblast | GSM3081987 |
| RNAseq-EryP-Rep3 | RNA-seq | Primitive erythroblast | GSM3081988 |
| RNAseq-EryD-Rep1 | RNA-seq | Definitive erythroblast | GSM3081989 |
| RNAseq-EryD-Rep2 | RNA-seq | Definitive erythroblast | GSM3081990 |
| RNAseq-EryD-Rep3 | RNA-seq | Definitive erythroblast | GSM3081991 |
| ATACseq-EryP_Rep1 | ATAC-seq | Primitive erythroblast | GSM3081992 |
| ATACseq-EryP_Rep2 | ATAC-seq | Primitive erythroblast | |
| ATACseq-EryD-Rep1 | ATAC-seq | Definitive erythroblast | GSM3081993 |
| ATACseq-EryD-Rep2 | ATAC-seq | Definitive erythroblast | |
| H3K27ac-EryP-Rep1 | ChIP-seq | Primitive erythroblast | GSM3081994 |
| H3K27ac-EryP-Rep2 | ChIP-seq | Primitive erythroblast | GSM3081995 |
| H3K27ac-EryD-Rep1 | ChIP-seq | Definitive erythroblast | GSM3081996 |
| H3K27ac-EryD-Rep2 | ChIP-seq | Definitive erythroblast | GSM3081997 |
| H3K4me1-EryP-Rep1 | ChIP-seq | Primitive erythroblast | GSM3081998 |
| H3K4me1-EryP-Rep2 | ChIP-seq | Primitive erythroblast | GSM3081999 |
| H3K4me1-EryD-Rep1 | ChIP-seq | Definitive erythroblast | GSM3082000 |
| H3K4me1-EryD-Rep2 | ChIP-seq | Definitive erythroblast | GSM3082001 |
| Gata1-EryP | ChIP-seq | Primitive erythroblast | GSM3082018 |
| Gata1-EryD | ChIP-seq | Definitive erythroblast | GSM3082019 |
| Tal1-EryP | ChIP-seq | Primitive erythroblast | GSM3082020 |
| Tal1-EryD | ChIP-seq | Definitive erythroblast | GSM3082021 |
| Input-EryP-Rep1 | ChIP-seq | Primitive erythroblast | GSM3082014 |
| Input-EryP-Rep2 | ChIP-seq | Primitive erythroblast | GSM3082015 |
| Input-EryD-Rep1 | ChIP-seq | Definitive erythroblast | GSM3082016 |
| Input-EryD-Rep2 | ChIP-seq | Definitive erythroblast | GSM3082017 |
| Myb-FL-Rep1 | ChIP-seq | E13.5 Fetal liver | GSM3082024 |
| Myb-FL-Rep2 | ChIP-seq | E13.5 Fetal liver | GSM3082025 |

| | | | |
|---|---|---|---|
| Input-FL | ChIP-seq | E13.5 Fetal liver | GSM3082026 |
| P300-FL-Fbbio-Rep1 | ChIP-seq | Fetal liver, P300-FB-biotin | GSM3082027 |
| P300-FL-Fbbio-Rep2 | ChIP-seq | Fetal liver, P300-FB-biotin | GSM3082028 |
| P300-FL-Fbbio-Rep3 | ChIP-seq | Fetal liver, P300-FB-biotin | GSM3082029 |
| Input-FL-Fbbio | ChIP-seq | Fetal liver, P300-FB-biotin | GSM3082030 |
| Gata1-Mel-CtrlshRNA | ChIP-seq | Mel cell, control shRNA | GSM3082031 |
| Gata1-Mel-MybshRNA | ChIP-seq | Mel cell, Myb shRNA | GSM3082032 |
| H3K27ac-Mel-CtrlshRNA | ChIP-seq | Mel cell, control shRNA | GSM3082033 |
| H3K27ac-Mel-MybshRNA | ChIP-seq | Mel cell, Myb shRNA | GSM3082034 |
| Input-Mel-CtrlshRNA | ChIP-seq | Mel cell, control shRNA | GSM3082035 |
| Input-Mel-MybshRNA | ChIP-seq | Mel cell, Myb shRNA | GSM3082036 |
| HiChIP-Gata1-EryP-Rep1 | HiChIP | E10.5 peripheral blood | |
| HiChIP-Gata1-EryP-Rep2 | HiChIP | E10.5 peripheral blood | |
| HiChIP-Gata1-EryP-Rep1 | HiChIP | E13.5 Fetal liver | |
| HiChIP-Gata1-EryP-Rep2 | HiChIP | E13.5 Fetal liver | |

**Supplemental Table S2: Total reads and valid peak numbers of ATAC-seq, ChIP-seq and HiChIP datasets.**

| Dataset | Total reads | Valid reads/ Valid pairs | Peaks |
|---|---|---|---|
| ATACseq-EryP_Rep1 | 39,716,421 | 14,528,709 | 52,193 |
| ATACseq-EryP_Rep2 | 33,903,736 | 16,997,888 | 43,843 |
| ATACseq-EryD-Rep1 | 32,379,234 | 12,480,432 | 33,510 |
| ATACseq-EryD-Rep2 | 101,177,115 | 95,831,929 | 56,178 |
| H3K27ac-EryP-Rep1 | 29,451,920 | 7,979,221 | 12,021 |
| H3K27ac-EryP-Rep2 | 32,800,601 | 11,384,609 | 16,117 |
| H3K27ac-EryD-Rep1 | 28,078,992 | 18,334,270 | 15,925 |
| H3K27ac-EryD-Rep2 | 22,102,546 | 12,011,039 | 17,363 |
| H3K4me1-EryP-Rep1 | 26,839,740 | 9,404,217 | 29,446 |
| H3K4me1-EryP-Rep2 | 18,154,855 | 8,407,207 | 39,359 |
| H3K4me1-EryD-Rep1 | 21,747,444 | 10,285,818 | 32,173 |
| H3K4me1-EryD-Rep2 | 24,235,323 | 8,931,552 | 24,940 |
| Gata1-EryP | 49,613,756 | 27,837,204 | 31,240 |
| Gata1-EryD | 50,693,265 | 34,139,464 | 30,324 |
| Tal1-EryP | 58,147,679 | 41,615,675 | 7,722 |
| Tal1-EryD | 56,987,926 | 39,583,235 | 9,621 |
| Input-EryP-Rep1 | 18,979,402 | 8,426,670 | N/A |
| Input-EryP-Rep2 | 32,107,945 | 25,316,253 | N/A |
| Input-EryD-Rep1 | 15,528,104 | 12,566,930 | N/A |
| Input-EryD-Rep2 | 18,312,587 | 9,128,574 | N/A |
| Myb-FL-Rep1 | 46,857,352 | 46,096,044 | 1,074 |
| Myb-FL-Rep2 | 34,576,877 | 34,280,120 | 1,259 |
| Input-FL | 34,798,097 | 34,535,830 | N/A |
| P300-FL-Fbbio-Rep1 | 78,310,708 | 64,757,816 | 13,542 |
| P300-FL-Fbbio-Rep2 | 74,401,171 | 62,491,054 | 27,665 |
| P300-FL-Fbbio-Rep3 | 68,837,801 | 56,949,368 | 10,307 |
| Input-FL-Fbbio | 83,951,131 | 81,937,175 | N/A |

| | | | |
|---|---|---|---|
| Gata1-Mel-CtrlshRNA | 30,678,121 | 30,005,070 | 16,742 |
| Gata1-Mel-MybshRNA | 16,681,056 | 16,251,064 | 3,911 |
| H3K27ac-Mel-CtrlshRNA | 14,267,695 | 13,952,508 | 28,157 |
| H3K27ac-Mel-MybshRNA | 14,279,704 | 13,952,062 | 19,103 |
| Input-Mel-CtrlshRNA | 31,222,795 | 30,708,929 | N/A |
| Input-Mel-MybshRNA | 14,595,861 | 33,085,147 | N/A |
| HiChIP-Gata1-EryP-Rep1 | 219,367,612 | 74,422,919 | N/A |
| HiChIP-Gata1-EryP-Rep2 | 310,236,431 | 71,122,380 | N/A |
| HiChIP-Gata1-EryD-Rep1 | 263,949,802 | 73,950,957 | N/A |
| HiChIP-Gata1-EryD-Rep2 | 332,206,672 | 30,871,740 | N/A |

**Supplemental Table S3, related to Fig. 3: List of target genes and shRNAs sequences.**

| Gene Name | Gene ID | Accession # | shRNA sequence |
|---|---|---|---|
| *Myb* | 17863 | NM_001198914.1 | AGCATTATCAGTCCGTCCG |
| | | NM_010848.3 | GGAGACGCCTGCGAGAACA |
| | | | AACAGGTGCACTATCCCCA |

**Supplemental Table S4, related to Fig. 4: List of target enhancers and guide RNAs sequences**

| Enhancer | Genomic coordinates | 5' gRNA sequence | 3' gRNA sequence |
|---|---|---|---|
| E1 | chr12: 111517811-111518523 | TGAAACTCACTAAGTAGAGC | CCACACTTCAGGTGATGAGA |
| E2 | chr4: 46410632-46411247 | TTAAGTTCATCCGTTGTGTT | CAGACTACAGCCGCCGGCTC |

**Supplemental Table S5, related to Fig. 4: List of genotyping PCR primers**

| Name | Amplification region | PCR primer Forward | PCR primer Reverse |
|---|---|---|---|
| E1 | Flanking | GGTCAGGTTTAGCAGCAAGC | TTAAGTTCCAGGGCAACCAG |
| E1 | Internal | GCGGCCAGATGATAATGAAT | CACACAATAGAATACATGCGTACAA |
| E2 | Flanking | CCCATGGCTTTGTACATGCT | CATCATCGCATTCTGTTTGG |
| E2 | Internal | TTCATGCACACTTAGCAGCA | GAGCCTCTCTGGGCTATGAG |

**Supplemental Table S6, related to Fig. 1 and Fig. 4: qRT-PCR primers**

| Gene Name | Gene ID | Accession # | Forward primer | Reverse primer |
|---|---|---|---|---|
| *Hbb-b1* | 15129 | NM_001278161.1 | TTTAACGATGGCCTGAATCACTT | CAGCACAATCACGATCATATTGC |
| *Hbb-bh1* | 15132 | NM_008219.3 | TGGACAACCTCAAGGAGACC | ACCTCTGGGGTGAATTCCTT |
| *Hbb-y* | 15135 | NM_008221.4 | TGGCCTGTGGAGTAAGGTCAA | GAAGCAGAGGACAAGTTCCCA |
| *Bcl11a* | 14025 | NM_016707.3 | AACCCCAGCACTTAAGCAAA | ACAGGTGAGAAGGTCGTGGT |
| *Gata1* | 14460 | NM_008089.2 | TGTCCTCACCATCAGATTCCA | TCCCTCCATACTGTTGAGCAG |
| *Tal1* | 21349 | NM_011527.3 | CGGCAGCAGAATGTGAATGG | CTCCTGGTCATTGAGTAACTTGG |
| *Trmo* | 74753 | NM_029086.2 | GAGCCAATAGGGTACTTGGAATC | CATCAAGGAATGCTCAGGGTTAT |
| *Hemgn* | 93966 | NM_053149.2 | CGACCTCGTCTGAAGCTCC | TCCTGTTCTCTGATACTTGCGT |
| *Anp32b* | 67628 | NM_130889.2 | AGCCGTTCGAGAACTTGTCTT | CAGGTTATTGCCACTTAGGTTCA |
| *Nans* | 94181 | NM_053179.3 | CAGAACCACCAAGGAGACATAGA | GGCCTTCCGGTTAAACTTGAAC |
| *Tnfaip2* | 21928 | NM_009396.2 | AGGAGGAGTCTGCGAAGAAGA | GGCAGTGGACCATCTAACTCG |
| *Eif5* | 217869 | NM_173363.5 | AAATCAGTGACCATGCAAAAGGT | TGCCTCAGCCACAATTTCTTTA |
